# Nested oscillatory dynamics in cortical organoids model early human brain network development

**DOI:** 10.1101/358622

**Authors:** Cleber A. Trujillo, Richard Gao, Priscilla D. Negraes, Isaac A. Chaim, Alain Domissy, Matthieu Vandenberghe, Anna Devor, Gene W. Yeo, Bradley Voytek, Alysson R. Muotri

**Affiliations:** University of California San Diego, School of Medicine, Department of Pediatrics/Rady Children’s Hospital San Diego, La Jolla, California 92093, USA.; University of California San Diego, School of Medicine, Department of Cellular & Molecular Medicine, La Jolla, California 92093, USA.; University of California San Diego, Department of Cognitive Science, Neurosciences Graduate Program, Institute for Neural Computation, La Jolla, California 92093, USA.; University of California San Diego, Department of Radiology, Department of Neurosciences, La Jolla, California 92093, USA; Harvard Medical School, Massachusetts General Hospital, Martinos Center for Biomedical Imaging, Charlestown, MA 02129, USA.; Molecular Engineering Laboratory, Agency for Science, Technology and Research, Singapore, Singapore.; Department of Physiology, Yong Loo Lin School of Medicine, National University of Singapore, Singapore, Singapore.; University of California San Diego, Kavli Institute for Brain and Mind, La Jolla, California 92093, USA.; Center for Academic Research and Training in Anthropogeny (CARTA), La Jolla, California 92093, USA.

**Keywords:** brain organoids, network oscillations, stem cells, phase-amplitude coupling, preterm electroencephalography (EEG), Methyl-CpG-binding protein 2 (MECP2)

## Abstract

Structural and transcriptional changes during early brain maturation follow fixed developmental programs defined by genetics. However, whether this is true for functional network activity remains unknown, primarily due to experimental inaccessibility of the initial stages of the living human brain. Here, we developed cortical organoids that spontaneously display periodic and regular oscillatory network events that are dependent on glutamatergic and GABAergic signaling. These nested oscillations exhibit cross-frequency coupling, proposed to coordinate neuronal computation and communication. As evidence of potential network maturation, oscillatory activity subsequently transitioned to more spatiotemporally irregular patterns, capturing features observed in preterm human electroencephalography (EEG). These results show that the development of structured network activity in the human neocortex may follow stable genetic programming, even in the absence of external or subcortical inputs. Our approach provides novel opportunities for investigating and manipulating the role of network activity in the developing human cortex.

**HIGHLIGHTS:**

- Early development of human functional neural networks and oscillatory activity can be modeled *in vitro*.
- Cortical organoids exhibit phase-amplitude coupling between delta oscillation (2 Hz) and high-frequency activity (100-400 Hz) during network-synchronous events.
- Differential role of glutamate and GABA in initiating and maintaining oscillatory network activity.
- Developmental impairment of MECP2-KO cortical organoids impacts the emergence of oscillatory activity.
- Cortical organoid network electrophysiological signatures correlate with human preterm neonatal EEG features.

**eTOC:** Brain oscillations are a candidate mechanism for how neural populations are temporally organized to instantiate cognition and behavior. Cortical organoids initially exhibit periodic and highly regular nested oscillatory network events that eventually transition to more spatiotemporally complex activity, capturing features of late-stage preterm infant electroencephalography. Functional neural circuitry in cortical organoids exhibits emergence and development of oscillatory network dynamics similar to those found in the developing human brain.

## INTRODUCTION

Diverse and hierarchical cellular networks develop into circuits with patterns of functional spatiotemporal activity to form the human brain. Neural oscillations, a prominent, rhythmic brain signal found across species, robustly track cognitive, behavioral, and disease states (Buzsáki and Draguhn, 2004; Fries, 2005; de Hemptinne et al., 2015; Henriques and Davidson, 1991; Khan et al., 2013; Uhlhaas and Singer, 2010), and have long been leveraged in cognitive and systems neuroscience due to their ubiquity and accessibility. These complex network dynamics emerge early in development, and is unclear if shaped exclusively by biological programming prenatally (Blankenship and Feller, 2010; Johnson, 2001; Power et al., 2010). *In vitro* and *in vivo* rodent studies have shown that a conserved repertoire of organized network activity, such as traveling waves, giant depolarizing potentials, and early network oscillations, develop according to a consistent timeline prior to and immediately after birth (Allene et al., 2008; Khazipov and Luhmann, 2006; Uhlhaas et al., 2010). However, due to an inability to interrogate the electrophysiology of intact embryonic brains, it remains unknown whether the same happens in humans. As a result, our knowledge about human brain functional development rests upon observations from nonhuman model systems (Power et al., 2010).

Organoids generated from induced pluripotent stem cells (iPSC) have emerged as a scaled-down and three-dimensional model of the human brain, mimicking various developmental features at the cellular and molecular levels (Camp et al., 2015; Lancaster and Knoblich, 2014; Lancaster et al., 2013; van de Leemput et al., 2014; Luo et al., 2016; Mariani et al., 2012; Paşca et al., 2015; Qian et al., 2016; Renner et al., 2017). Despite recent advances in the understanding of their vast cellular diversity, there is no evidence that these organoids develop complex and functional neural network activity that resembles early human brain formation (Birey et al., 2017; Quadrato et al., 2017). Therefore, researchers have not yet clearly determined whether organoids are a suitable model for neural network dynamics (Kelava and Lancaster, 2016; Paşca, 2018).

Here, we use human iPSCs to generate cortical organoids that exhibit evolving and nested oscillatory network dynamics over the span of several months. We subsequently investigated the molecular basis of human brain oscillatory activity formation, maintenance, and temporal control by gene targeting. Finally, we applied unsupervised machine learning to evaluate the similarity between electrophysiological activity patterns of the *in vitro* model and human preterm neonatal electroencephalogram (EEG). Our findings suggest that organoid models are suitable for the investigation of the physiological basis of network formation at early and late stages of the human brain development. This prolonged evaluation of cortical organoid activity expands our understanding of the emergence of network-level neurodynamics in humans.

## RESULTS

### Generation of functional cortical organoids

Despite the structural and transcriptional similarities between brain organoids and the developing nervous system, the emergence of higher-level complex network activity comparable to the living human brain remains largely untested (Figure 1A). To investigate the formation of a functional network, we promoted cortical specification by modifying previously described protocols (Paşca et al., 2015; Thomas et al., 2016) (Figure 1B, see Methods for details). At the beginning of differentiation, an abundance of proliferative neural progenitor cells (NPCs) (Ki67+, SOX2+ and Nestin+) that self-organized into a polarized neuroepithelium-like structure was observed. Similar to human cortical development *in vivo*, the proliferative zone around a lumen delimited by β-catenin+ cells was surrounded by progenitor cells. Progressively, the organoids increased in size and in the proportion of mature neurons (NeuN+ and MAP2+) to ultimately develop into concentric multi-layer structures composed of NPCs, intermediate progenitors (TBR2+, also known as EOMES), and lower (CTIP2+, also known as BCL11B) and upper (SATB2+) cortical layer neurons (Figure 1B-E and S1A-C). Although the initial fraction of glial cells was less than 5%, this population increased to about 30-40% after 6 months of differentiation (Figure 1D, 1E and S1D-H). The neurons exhibit dendritic protrusions and synaptic structures (Figure 1F and 1G).

**Figure 1.**
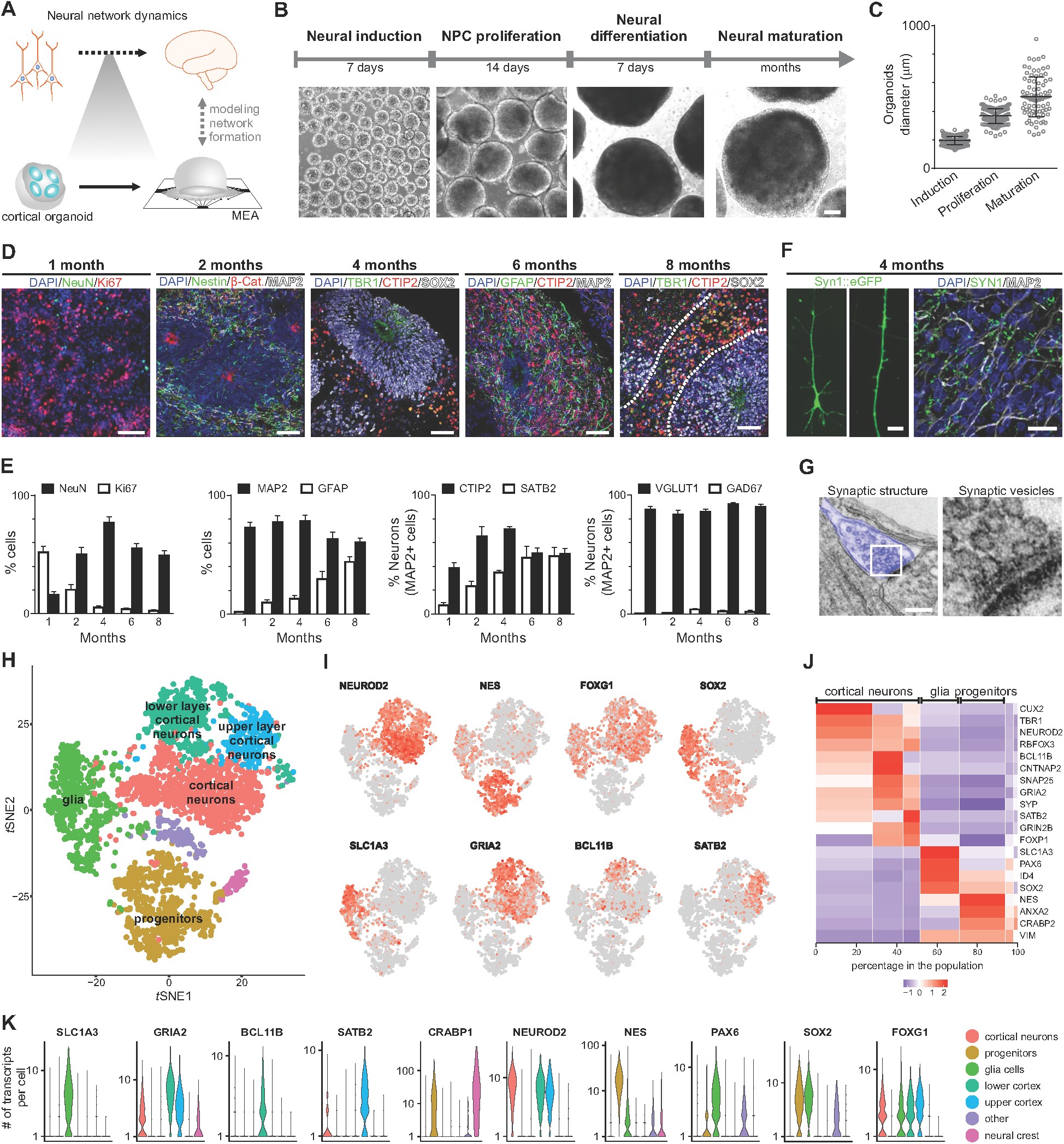
Cellular and molecular development of human cortical organoids. (A) Overview of human neural network formation and dynamics evaluation using organoids. (B) Schematic of the protocol used to generate cortical organoids. Scale bar, 200 μm. (C) Organoid growth during different developmental stages. (D) Representative immunostainings showing proliferating NPCs (Ki67+ and Nestin+), lower (TBR1+ and CTIP2+) and upper (SATB2+) cortical layer neurons and glial cells (GFAP+) overtime. Scale bar, 50 μm. (E) Population analysis of specific markers indicating stages of maturation and multiple neuronal subtypes. The data are shown as mean ± s.e.m. (*n = 8*). (F) Representative image of a pyramidal neuron (left panel); dendritic structures are observed in cells transduced with the SYN:EGFP reporter (middle panel; scale bar, 5 μm). Immunohistochemical detection of the synaptic protein Syn1 (right panel; scale bar, 50 μm). (G) Electron microscopy of synaptic structures in 4-month-old cortical organoids (blue). (H) *t*-distributed stochastic neighbor embedding (*t*SNE) plot of 3,491 cells from 6-month-old organoids. Colors denote seven main cell clusters. (I) *t*SNE plots depicting cell-type specific marker expression levels (red denotes higher expression). (J) Heatmap of average expression for representative gene markers by cluster and cell-type (see also Figure S4). (K) Violin plots showing transcript levels for representative markers of each cluster (see Figure S3 for additional markers).

To characterize the cellular diversity of a cortical organoid, we performed single-cell gene expression profiling in 6-month-old organoids and used unbiased clustering to classify the main existing cell types. From two independent differentiation replicates (Figure S2A-D), seven distinct clusters were characterized based on their differential gene expression patterns including: progenitors, glia, and cortical neurons, which could be further subdivided into lower and upper layer based on the expression of the layer-specific markers CTIP2 and SATB2, respectively (Figure 1H-K, S2E, S2F and Table S1).

### Emergence of nested oscillatory network activity

In addition to the observed cellular diversity and expression of synaptic markers, we interrogated the presence of functional network activity. We performed weekly extracellular recordings of spontaneous electrical activity using multi-electrode arrays (MEA). Six-week-old cortical organoids were plated per well in 12-well MEA plates contains 64 platinum microelectrodes with 30 μm of diameter spaced by 200 μm, yielding a total of 512 channels. We separately analyzed single-channel and population firing characteristics derived from channel-wise spike times, and the local field potential (LFP); a measure of aggregate synaptic currents and other slow ionic exchanges (Buzsáki et al., 2012) (Figure 2A). The spikes from each channel do not represent putative single-unit action potentials. Since the spatial resolution of MEA electrodes was sparse, the total population spiking of a well was submitted for further analysis, rather than individual spike trains. Over the course of 10 months, organoids exhibited consistent increases in electrical activity, as parametrized by channel-wise firing rate, burst frequency, and spike synchrony (Figure 2B-D and S3A-E), which indicates a continually-maturing neural network (Chen et al., 2009; Lisman, 1997). Additionally, the variability between replicates over 40 weeks of differentiation was significantly lower compared to iPSC-derived neurons in monolayer cultures (Figure 2C inset and S3E).

**Figure 2.**
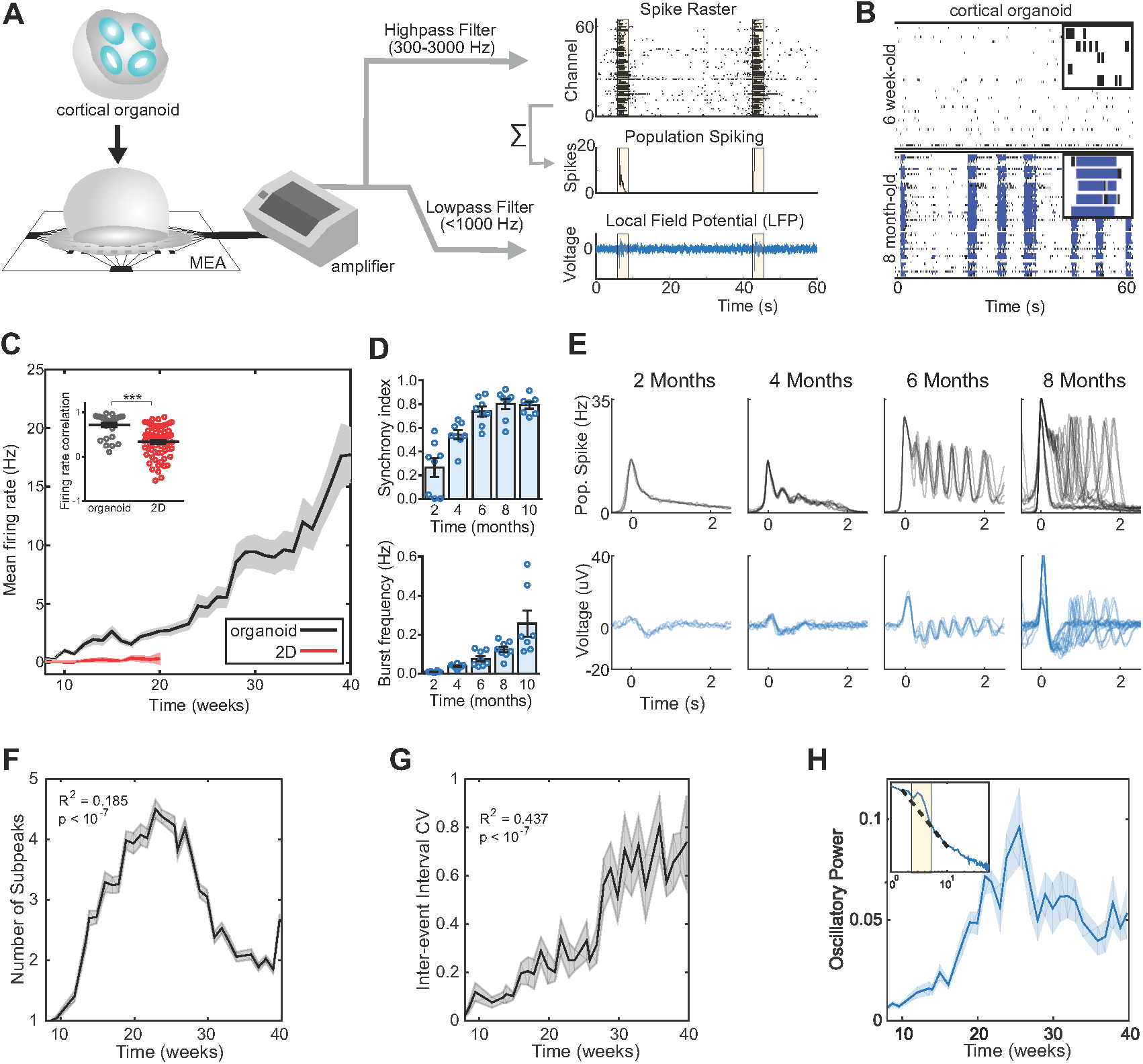
Oscillatory network dynamics in long-term cortical organoids. (A) Schematic of the organoid signal processing pipeline. Raw MEA data is analyzed as population spiking and LFP separately. Synchronous network events are highlighted in yellow. (B) Raster plot of network spiking activity after 1.5 and 6 months of maturation. A 3-s interval of activity over 5 channels is shown in the upper right corners. (C) Cortical organoids show elevated and continuously increasing mean firing rate compared to 2D monolayer neurons (*n = 8* organoid cultures, and *n = 12* for 2D neurons). Inset, correlation of the firing rate vector over 12 weeks of differentiation (from 8 to 20) between pairs of cultures showing reduced variability among organoid replicates. (D) Temporal evolution of cortical organoid network activity. Detailed definitions and further parameters are presented in Figure 5B and 5C. (E) Time series of population spiking and LFP during network events in cortical organoid development. Each trace represents a single event during the same recording session. (F) Oscillatory dynamics within network events develop nonlinearly, following an inverted-U trajectory. (G) Increase of network variability dynamics throughout development. (H) Oscillatory power increases up to the 25th week in culture and plateaus at 30 weeks. Inset, Oscillatory power is calculated by fitting a straight line (dashed) over the aperiodic portion of the PSD and taken as the height of narrow peaks rising above the linear fit. The data shown in C, D, F, G and H are presented as mean ± s.e.m. *P < 0.05, **P < 0.01, ***P < 0.001, unpaired Student’s *t*-test (C), quadratic (F) and linear (G) regression.

During individual recordings, cultures displayed a robust pattern of activity, switching between long periods of quiescence and short bursts of spontaneous network-synchronized spiking (hereafter referred to as “network events”). These network events are periodic (∼0.05 Hz) but infrequent early in development (∼2 months), occurring roughly every 20 seconds and decayed monotonically after the initial onset, similar to previously reported network “oscillations” in primary cultures and organoids (Figure 2E). From 4-months onwards, a secondary peak emerged 300-500 ms after the initial network activation, leading to the presence of a nested fast oscillatory (2-3 Hz) pattern up to 6-months in culture (Figure 2F and Figure S4A-G). Notably, this robust fast timescale nested oscillation was not observed in 3D neurospheres, suggesting that the spherical arrangement of neurons is insufficient for the emergence of nested oscillations (Figure S4H-J). The regular oscillatory activity during network events transitioned to stronger, yet more variable, oscillations over time. To quantify this network complexity, we tracked the regularity (coefficient of variation of inter-event intervals, CV) and the spatial and temporal correlation between spontaneous network events. The inter-event interval CV consistently increased over 10 months of differentiation (Figure 2G), from extremely regular latencies (CV ≅ 0) at 2 months to irregular, Poisson-like (CV ≅ 1) at 10 months. This indicates increased variability between consecutive network events initiation. Additionally, spatial and temporal irregularity on a shorter time-scale (within-event) also increased with development, suggesting a breakdown of deterministic population dynamics from the onset of network events (Figure S4G).

Periodic oscillatory activity is often defined as a “bump” over the characteristic *1/f* background noise in the power spectral density (PSD) of extracellular signals above-and-beyond the aperiodic *1/f* signal (Buzsáki et al., 2013; Gao et al., 2017). In organoid LFPs, we observed both prominent oscillatory peaks in the low-frequency range (1-4 Hz) and in the aperiodic signal characteristic of neural recordings (Ben-Ari, 2001; Voytek et al., 2015). The development of oscillatory activity in cortical organoids over time was quantified by computing the PSD for each LFP recording (Figure 2H, inset). Oscillatory power in the delta range (1-4 Hz) increased for up to 24 weeks in culture, tapering off slightly in subsequent recordings and plateauing during the last 10 weeks. This inverted-U trajectory reflects the network’s initial acquisition of oscillatory modes at steady frequencies and the dispersion of this regularity at later time points. The LFP results reveal the development of the cortical organoid cultures across different network states: from sparse activity with extreme rigidity and regularity, to one that acquires repetitive, perhaps overly-regular oscillatory patterns (Voytek and Knight, 2015), until it finally reaches a stage of higher spatiotemporal complexity and variability that is reminiscent of self-organized networks (Tetzlaff et al., 2010) (Figure S4C-G).

### Oscillatory coordination of neural ensembles and its synaptic mechanisms

Oscillatory dynamics in the functioning brain have been postulated to coordinate spiking across neural ensembles. In the LFP and other mesoscopic brain signals, this manifests as a phenomenon known as cross-frequency phase-amplitude coupling (PAC) (Voytek and Knight, 2015), wherein the high-frequency content of the LFP is entrained to the phase of slow oscillations (Manning et al., 2009; Miller et al., 2007; Mukamel et al., 2005). PAC in the neocortex and hippocampus has been shown to be functionally relevant in a range of behaviors and neurological disorders (de Hemptinne et al., 2015; Voytek and Knight, 2015; Voytek et al., 2015). In the organoids, we observed greater PAC between oscillatory delta (1-4 Hz) and broadband gamma activity (100-400 Hz, see Methods) during network events compared to quiescent periods (Figure 3A-C).

**Figure 3.**
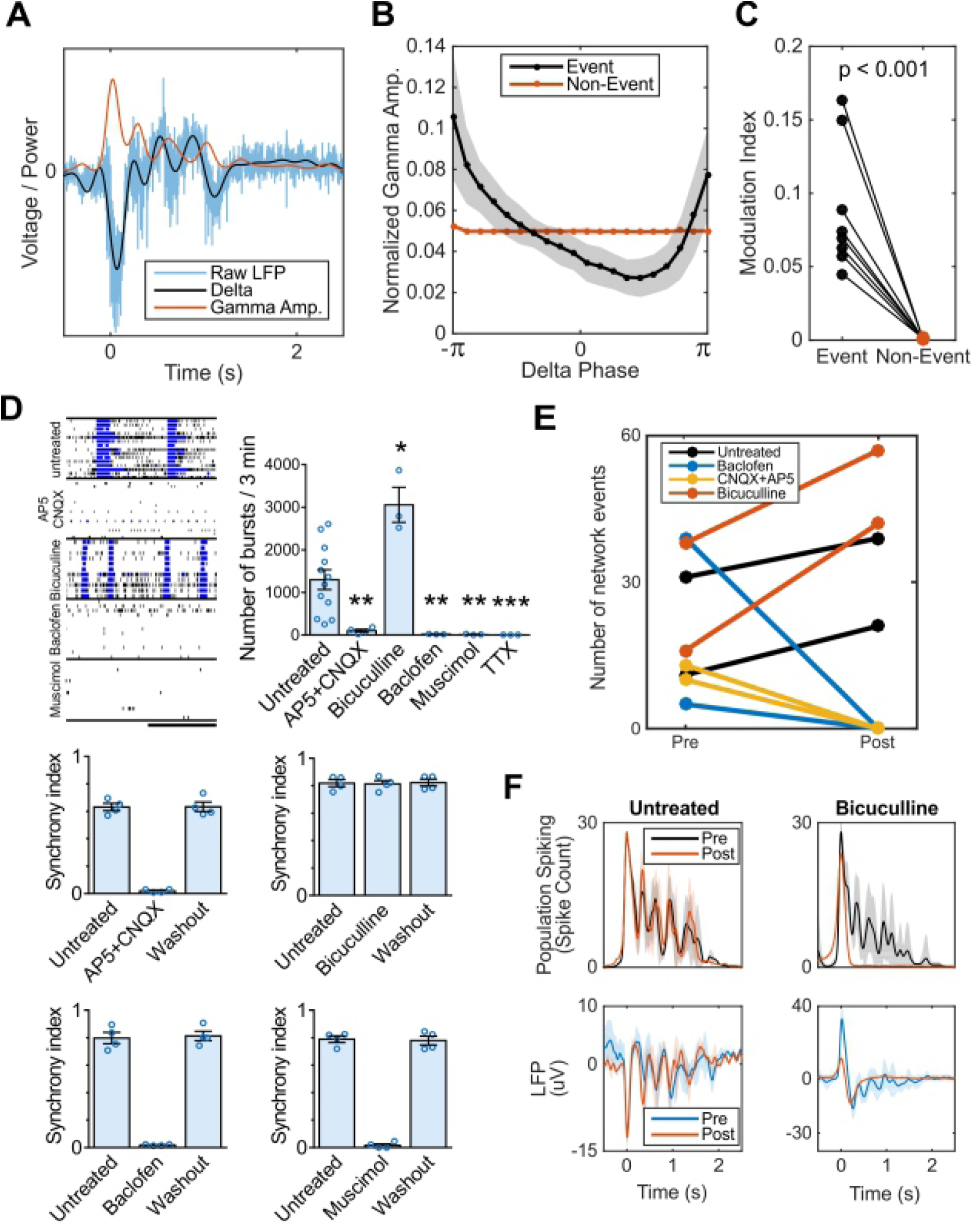
Cortical organoid serves as a model of functional oscillations and their synaptic mechanisms. (A-C) Phase-amplitude coupling is observed in organoid LFP during network events, a phenomenon proposed to mediate neural communication *in vivo*. (A) Example of raw LFP during a network event decomposed into its low-frequency component (1-4 Hz delta) and the amplitude envelope of the high-frequency, broadband gamma component (200-400 Hz). Analysis was repeated for 100-200 Hz with near identical effect size and significance. (B) Normalized gamma amplitude binned by delta phase during network events (black) shows greater modulation depth by low frequency delta than during non-event periods (red). (C) Phase-amplitude coupling during network events is significantly greater than non-event periods in all batches. (D) Effect of selective drug treatments on neuronal electrical activity in 6-month-old organoids. Representative raster plots and burst measurements of untreated and treated organoids. Scale bar, 20 s. Exposure to AP5 + CNQX, baclofen and muscimol reversibly extinguish the network bursts (synchrony), while no changes were promoted by bicuculline. (E-F) Pharmacological perturbation of oscillatory activity during network events in 6-month-old organoids. Application of bicuculline increases the number of network events, while CNQX + AP5 and baclofen completely abolish synchronized network events. Bicuculline blocks oscillatory network activity but not the network event itself. Data are shown as mean ± s.e.m.; unpaired Student’s *t*-test.

We further evaluated the role of glutamatergic and GABAergic synaptic transmission in forming oscillations by pharmacological intervention. Organoid neural networks were susceptible to both glutamate receptor antagonists (AP5 and CNQX; NMDA and AMPA/kainate, respectively) and GABA receptor agonists (muscimol, GABA_A_; baclofen, GABA_B_) by significantly reducing the number of spikes and bursts, with a subsequent extinction of synchronous activity. The electrical activity was abolished in the presence of tetrodotoxin (TTX) (Figure 3D and 3E). Notably, blockade of GABAergic transmission by bicuculline increased the number of network-synchronized events and did not affect peak population firing rates, but abolished nested 2 Hz oscillatory activity by erasing subsequent reverberant peaks (Figure 3F). The findings suggest that GABA transmission is crucial for the maintenance, but not the initiation of faster oscillatory activity. This is consistent with accounts of inhibition rhythmically coordinating pyramidal populations activity during early development (Opitz et al., 2002).

### MECP2 is essential for the timely emergence of network oscillations

In addition to modeling the typically-developing brain, cortical organoids can also shed light on the mechanism behind functional deficits in neurodevelopmental disorders (Birey et al., 2017; Lancaster et al., 2013; Thomas et al., 2016). Normal nested oscillatory network dynamics in the brain are often shown to break down in psychiatric and neurological conditions (Uhlhaas and Singer, 2010). However, the mechanisms by which that happens and its impact on the circuit are difficult to elucidate. Thus, we next investigated whether cortical organoids could be used to model oscillatory network defects. Previous work evidenced that patients with autism spectrum disorder exhibit reduced alpha oscillation power (8-12 Hz) and evoked low-gamma (40-60 Hz) response, as well as reduced PAC (Khan et al., 2013; Mohammad-Rezazadeh et al., 2016). Mutations in the Methyl-CpG-binding protein 2 (*MECP2*) gene lead to a severe disruption in cortical development that account for many symptoms of Rett syndrome, autism, schizophrenia and other neurological disorders (Amir et al., 1999; Cohen et al., 2002; Du et al., 2016; Liu et al., 2016; Wen et al., 2017). MECP2 is involved on the epigenetic regulation of target genes by binding to methylated CpG dinucleotides promoter regions, acting as a transcriptional modulator (Figure 4A).

**Figure 4.**
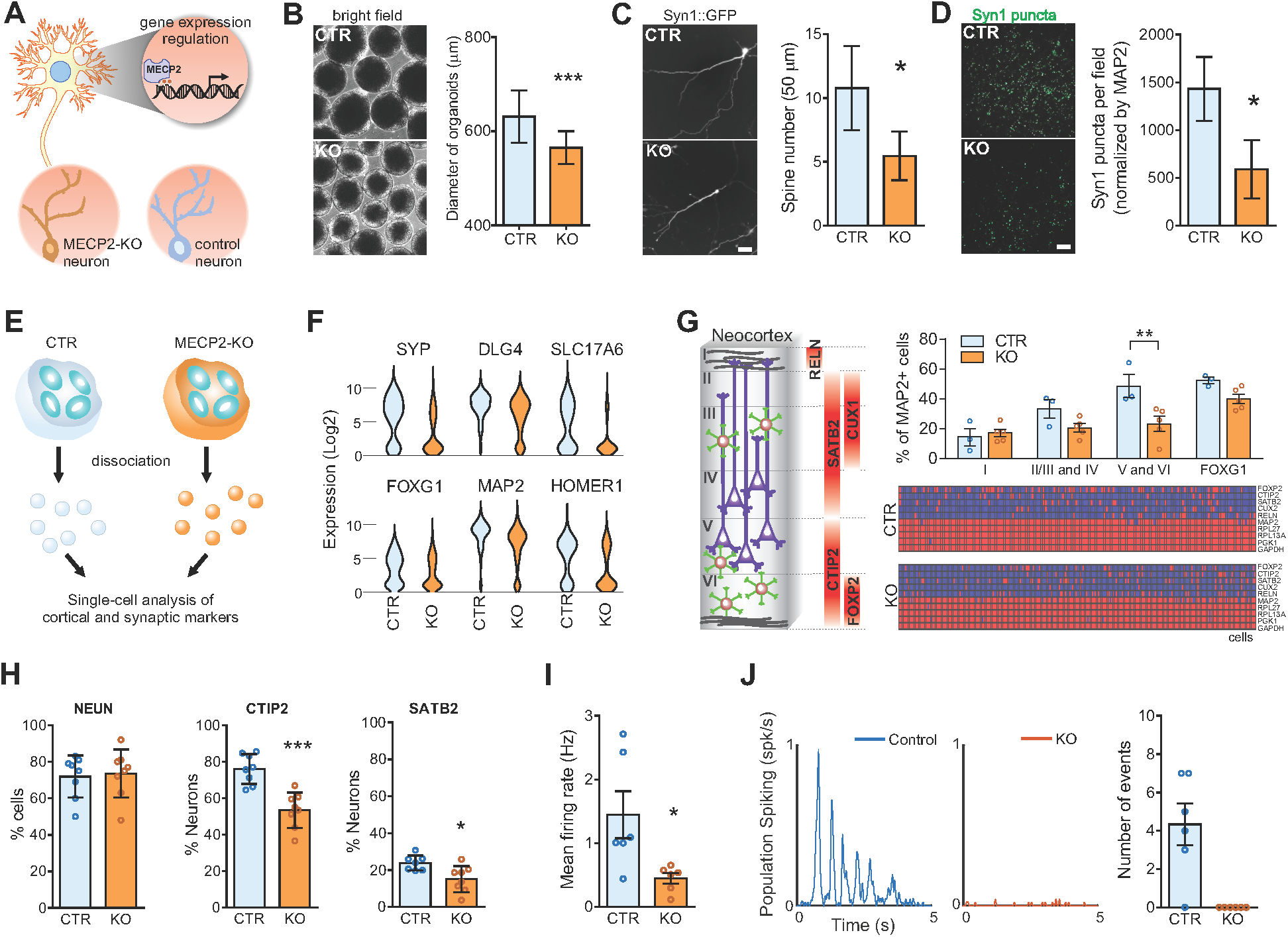
MECP2 contribution to the emergence of network oscillations. (A) MECP2-knockout neurons (MECP2-KO) show reduced spine-like density and soma size compared to controls. (B) Organoid diameter quantification (CTR, *n = 210* organoids; KO, *n = 333* organoids). (C) Spine-like density and (D) synaptic puncta are reduced in MECP2-KO neurons. Scale bar, 50 μm. (E-H) Targeted single-cell analysis of neural markers and cortical layer-related genes over defined control *Ct* value. In 3-month-old cortical organoids, a significant decrease in the number of CTIP2+ and SATB2+ neurons was observed. (I) MECP2-KO cortical organoids show decreased mean firing rate after 5 months of maturation (*n = 6* organoid cultures). (J) Lack of oscillatory network events in 5-month-old MECP2-KO organoids. Each trace represents a single event during the same recording session. For B, C, D, G, H, I and J, data are shown as mean ± s.e.m.; *P < 0.05, **P < 0.01, ***P < 0.001, unpaired Student’s *t*-test.

To model MECP2 deficiency during neurodevelopment, we generated a pluripotent stem cell model from two different cell lines, each carrying a distinct *MECP2* mutation that results in a nonfunctional protein (Zhang et al., 2016). Human *MECP2*-mutant neurons *in vitro* exhibit fewer synapses, smaller soma size, altered calcium signaling and electrophysiological defects compared to controls (Marchetto et al., 2010). Based on the observed reduction in the number of layer V neurons in *Mecp2*-mutant mice (Stuss et al., 2012) and documented clinical data of microcephaly in Rett syndrome patients (Amir et al., 1999), we sought to examine transcriptomics, cellular and structural differences using MECP2-KO cortical organoids. We observed a significant decrease in the diameter of MECP2-KO organoids, neuronal protrusions or spine-like density and synaptic puncta at later stages of differentiation (Figure 4B-D). Additionally, and similar to the *Mecp2*-mutant mice (Stuss et al., 2012), a significant reduction in the proportion of CTIP2+ and SATB2+ neurons was observed by targeted single-cell analysis (Figure 4E-G) and corroborated by immunostaining (Figure 4H). MECP2-KO cortical organoids also showed reduced neural activity leading to an absence of network oscillations when compared to isogenic control organoids at the same age (Figure 4I and 4J). The inability to entrain into a functionally connected network at early stages of development might underlie the core deficits found in MECP2-deficient related disorders. More importantly, these results highlight the contribution of specific genes in the timely emergence of oscillatory activity.

### Organoid network development recapitulates preterm EEG

Despite emergence of complex oscillatory network activity in organoids, it is unclear whether the spontaneous developmental trajectory observed is representative of programmed early neurodevelopment. While network activity from organoids does not exhibit the full temporal complexity seen in adults, the pattern of alternating periods of quiescence and network-synchronized events resembles electrophysiological signatures present in preterm human infant EEG. During *trace discontinu* (Tolonen et al., 2007), quiescent periods are punctuated by high-amplitude oscillations (spontaneous activity transients, SATs) lasting a few seconds. Intervals of complete quiescence disappear as infants become of term, and the EEG is dominated by continuous and low-amplitude desynchronized activity in adult brains (Figure 5A and S5A).

**Figure 5.**
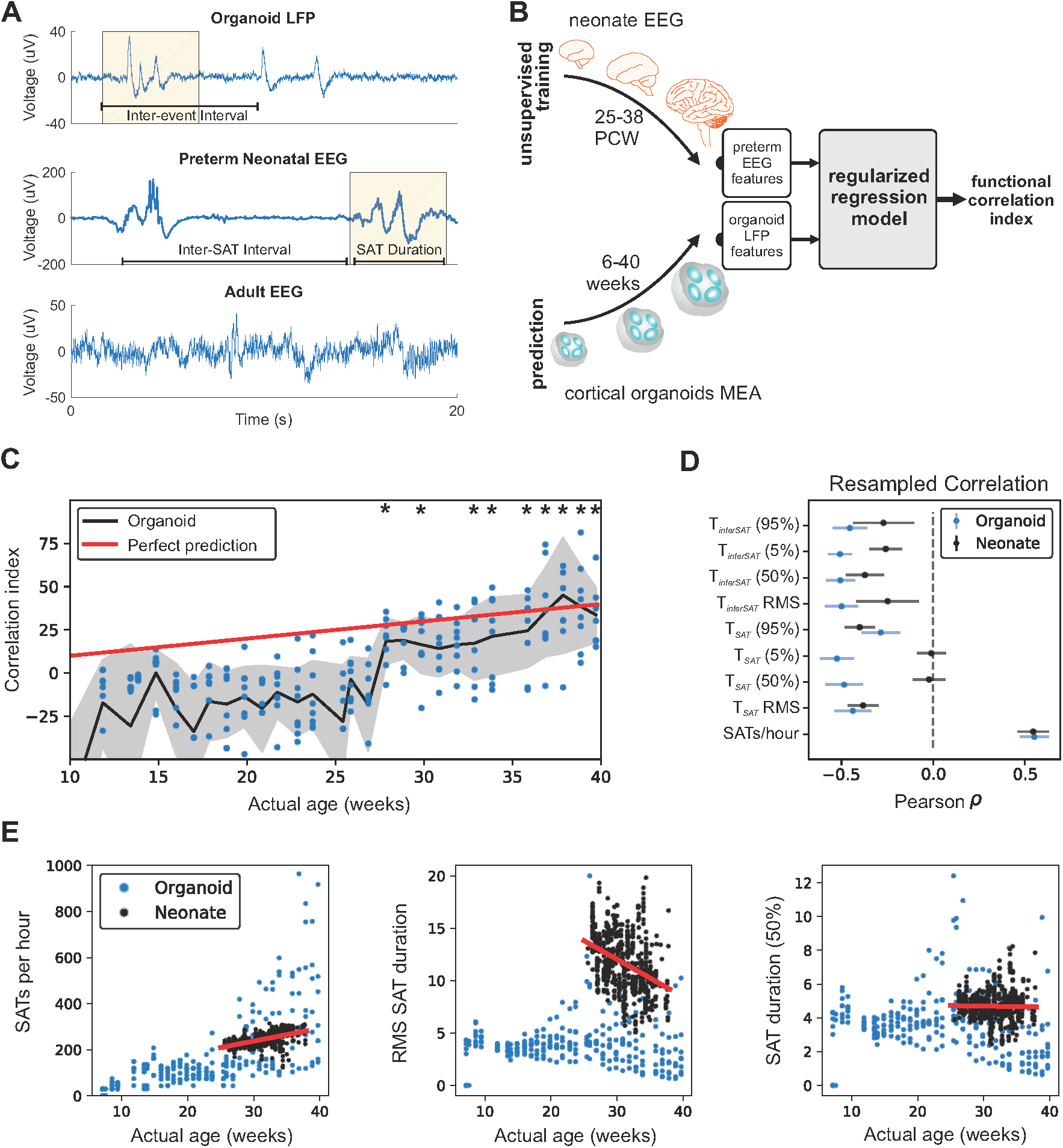
Organoid network dynamics mimic premature neonates after 28 weeks of maturation. (A) Representative LFP trace from cortical organoid, highlighting instances of network events (yellow). Comparable events between periods of quiescence (discontinuous network dynamics) are shown in human preterm neonate EEG at 35 weeks gestational age, while a different pattern of continuous activity is observed in adult EEG. SAT: spontaneous activity transient. (B) Schematic of unsupervised machine learning pipeline: 9 EEG features from 39 premature babies (*n* = 567 recordings) between 25 and 38 PCW were used to train and cross-validate a regularized regression model (ElasticNet) to first determine neonate brain age, which was then applied directly to organoid LFP features. (C) Machine-determined organoid “brain age” plotted against actual organoid age. Black stars denote time points where mean determined age is not significantly different from actual age under 1-sample *t*-test (P < 0.05, *n = 8*). (D) Resampled Pearson’s correlation coefficient between age and electrophysiological features for both organoid and premature neonates show different degrees of developmental similarity for individual features. (E) EEG/LFP features over time for organoids and premature neonates show various levels of similarity.

Because of the inability to interrogate the electrophysiology of intact human embryonic brains, we attempt to quantitatively compare network activity in cortical organoids to preterm human EEG, by applying an unsupervised regularized regression model (with cross-validation) (L1 & L2 regularized, ElasticNet) on a dataset of 567 preterm neonatal EEGs, ranging from 24 to 38 post-conception weeks (PCW) (Stevenson et al., 2017). Specifically, the training dataset consists of a subset of the preterm EEG data after we discarded features not sensibly computed from organoid LFPs, such as interhemispheric synchrony and frequency-dependent filtering properties of the skull (see Table S2 for a full list of included and rejected features). The remaining features correspond to aspects of spontaneous activity transient (SAT) timing, such as SATs per hour and SAT duration, which were similarly computed on organoid LFPs after network event detection.

Initially, the regression model was only optimized to anticipate preterm infant age based on their own brain features, and has not seen any organoid data whatsoever up to this point. Then, after unbiased training, we submitted analogous features computed from cortical organoid LFPs to the model for neurodevelopmental correlation (Figure 5B). Notably, brain organoids past 28 weeks in culture exhibit similar developmental trajectories of electrophysiological features as preterm neonates (Figure 5C). Next, we examined the similarities between brain organoids and preterm humans by looking at each specific feature (Figure 5C and 5D). Of all features, “SATs per hour” (“events per hour” in organoids) showed high similar values and growth, while “root-mean-square SAT duration” showed a similar decline (but not in absolute value) over 25 to 38 weeks in both datasets (Figure 5E, S5B and 5C). Therefore, while the developmental trajectory of cortical organoids is not identical to that of the fetal brain, a machine learning model trained only on preterm neonatal EEG features was able to demonstrate that the observed network electrophysiological features may share similarities representative of genetically programmed developmental timelines.

## DISCUSSION

Development of functional human brain networks is an activity-dependent process guided by genetic and molecular programs, shaped by the emerging cellular diversity. Neonatal neural networks share many features with adult brains, despite the fundamental structural differences (Power et al., 2010). Even though the chronological stages of the human cortical network formation are not well understood, it is suggested that emerging cognitive functions during infancy are a result of different brain regions and environmental cues (Johnson, 2001). However, *in utero* development is vital for the establishment of neuronal circuitry and healthy functioning of the brain. The second and third trimester of human gestation are when the corticothalamic network is formed *via* transient connections of the subplate GABAergic neurons and the emergence of synchronized network activity (Kostović and Judaš, 2010). Thus, early cortical functional maturation follows an independent sensory-input pathway, guided by spontaneous activity and associated with synaptic regulating mechanisms (Uhlhaas et al., 2010).

Here we report the formation of small-scale functional electrophysiological networks in human cortical organoids, with some similar features to those observed in the developing brain. While we do not claim functional equivalence between the organoids and a full neonatal cortex, the current results represent the first step towards an *in vitro* model that captures some of the complex spatiotemporal oscillatory dynamics of the human brain. Robust extracellular electrical activity was established at earlier stages and progressively developed into an organized oscillatory network. As such, we show that some features of early functional network dynamics (e.g., spontaneous activity transients) can be recapitulated by an *in vitro* model of the developing cortex, with no additional constraints other than structural and genetic similarities. This offers initial evidence for a convergent experience-independent neurodevelopmental program of the neocortex prior to birth. Given the potential roles of synchronized and oscillatory network dynamics in coordinating information flow between developed cortical brain regions during human cognition (Uhlhaas et al., 2010), these results highlight the potential for cortical organoids to advance our understanding of functional electrophysiology, brain development, and neuro-genetic disorders. The *MECP2* deficiency, leading to cellular and network defects, is an example of how this platform can be used to identify important genes that are essential for the timely emergence of oscillations in brain organoids. Finally, our findings may ultimately reframe the ethical discussions on human brain organoid research and offer an innovative link between microscale organoid physiology and cognitive neuroscience.

## SUPPLEMENTAL INFORMATION

Supplemental information includes Supplemental Experimental Procedures, 5 Supplemental Figures, 2 Supplemental Tables can be found with this article online.

## ACKNOWLEDGMENTS

This work was supported by grants from the California Institute for Regenerative Medicine (CIRM) DISC1-08825 and DISC2-09649, the National Institutes of Health through the R01MH108528, R01MH094753, R01MH109885, R01MH100175, R56MH109587, a SFARI grant #345469, and a NARSAD Independent Investigator Grant to A.R.M. B.V is supported by a Sloan Research Fellowship, the Whitehall Foundation (2017-12-73), and the National Science Foundation (1736028). I.A.C. is a San Diego IRACDA Fellow supported by National Institutes of Health (NIH)/NIGMS K12 GM06852 Award. G.W.Y. and A.R.M. are supported by U19MH1073671, part of the National Cooperative Reprogrammed Cell Research Groups (NCRCRG) to Study Mental Illness. R.G is supported by the Natural Sciences and Engineering Research Council of Canada (NSERC PGS-D), UCSD Kavli Innovative Research Grant (IRG), Frontiers for Innovation Scholars Program, and Katzin Prize. We thank Patrick S. Cooper for his help on the Australian EEG Database. We thank Dr. V.Taupin for her expert assistance with electron microscopy.

## AUTHOR CONTRIBUTIONS

C.A.T., R.G. and P.D.N. should be considered co-first authors as they each designed the experiments and conducted the analyses with input from A.R.M., B.V.; P.D.N. and C.A.T. generated and characterized the cortical organoids and performed the MEA recordings. C.A.T. performed C1 single-cell analyses and synaptic quantification. I.A.C. performed and analyzed 10X Genomics single-cell experiments. A.Do. processed 10X Genomics single-cell data. G.W.Y. led and funded the single-cell RNA-seq analyses. M.V. and A.De. performed the functional experiment. P.D.N. analyzed the MEA data using the Axion Biosystems Neural Metrics Tool. R.G. performed the custom MEA and EEG analyses. A.R.M. and B.V. should be considered co-senior authors as they contributed equally to directing the overall study design, with A.R.M. leading the cortical organoid development and analyses, and B.V. leading the electrophysiological design and analyses. C.A.T., R.G., P.D.N., B.V. and A.R.M wrote the manuscript. All authors reviewed the manuscript for publication.

## CONFLICT OF INTERESTS

Dr. Muotri is a co-founder and has equity interest in TISMOO, a company dedicated to genetic analysis focusing on therapeutic applications customized for autism spectrum disorder and other neurological disorders with genetic origins. The terms of this arrangement have been reviewed and approved by the University of California San Diego in accordance with its conflict of interest policies.

## AUTHOR INFORMATION

Correspondence and requests for materials should be addressed to

## EXPERIMENTAL PROCEDURES

### Cell source

iPSC lines derived from control individuals have been previously characterized elsewhere (Gore et al., 2011; Nageshappa et al., 2016). Human embryonic stem cell (ESC) and iPSC colonies were expanded on Matrigel-coated dishes (BD Biosciences, San Jose, CA, USA) with mTeSR1 medium (StemCell Technologies, Vancouver, Canada). The cells were routinely checked by karyotype and CNV arrays to avoid genomic alterations in the culture. The study was approved by the University of California San Diego IRB/ESCRO committee (protocol 141223ZF).

### MECP2-KO cell line generation

MECP2-deficient cell lines were generated by inducing pluripotency in fibroblasts derived from a male patient. Additionally, we used H9 human ESC with the CRISPR/Cas9 genome-editing system to induce frameshift mutations in the *MECP2* locus. This incorporation resulted in the creation of early stop codons rendering a non-functional MECP2 protein. Mutagenesis and off-targets were confirmed by exome sequencing techniques. The CRISPR-Cas protocol can be found elsewhere (Thomas et al., 2017). Once we confirmed the pluripotency state of the cellular models, we differentiated them into 2D neuronal monolayer cultures (Thanathom et al., 2016) and cortical organoids.

### Generation of cortical organoids

Feeder-free iPSCs were fed daily with mTeSR1 for 7 days. Colonies were dissociated using Accutase (Life Technologies, Carlsbad, CA, USA) in PBS (1:1) for 10 minutes at 37 °C and centrifuged for 3 minutes at 150 x *g*. The cell pellet was resuspended in mTeSR1 supplemented with 10 μM SB431542 (SB; Stemgent, Cambridge, MA, USA) and 1 μM Dorsomorphin (Dorso; R&D Systems, Minneapolis, MN, USA). Approximately 4 × 10^6^ cells were transferred to one well of a 6-well plate and kept in suspension under rotation (95 rpm) in the presence of 5 μM ROCK inhibitor (Y-27632; Calbiochem, Sigma-Aldrich, St. Louis, MO, USA) for 24 hours to form free-floating spheres. After 3 days, mTeSR1 was substituted by Media1 [Neurobasal (Life Technologies) supplemented with Glutamax, 2% Gem21 NeuroPlex (Gemini Bio-Products, West Sacramento, CA, USA), 1% N2 NeuroPlex (Gemini Bio-Products), 1% MEM nonessential amino acids (NEAA; Life Technologies), 1% penicillin/streptomycin (PS; Life Technologies), 10 μM SB and 1 μM Dorso] for 7 days. Then, the cells were maintained in Media2 [Neurobasal with Glutamax, 2% Gem21 NeuroPlex, 1% NEAA and 1% PS] supplemented with 20 ng/mL FGF2 (Life Technologies) for 7 days, followed by 7 additional days in Media2 supplemented with 20 ng/mL of FGF2 and 20 ng/mL EGF (PeproTech, Rocky Hill, NJ, USA). Next, cells were transferred to Media3 [Media2 supplemented with 10 ng/mL of BDNF, 10 ng/mL of GDNF, 10 ng/mL of NT-3 (all from PeproTech), 200 μM L-ascorbic acid and 1 mM dibutyryl-cAMP (Sigma-Aldrich)]. After 7 days, cortical organoids were maintained in Media2 for as long as needed, with media changes every 3-4 days.

### Mycoplasma testing

All cellular cultures were routinely tested for mycoplasma by PCR. Media supernatants (with no antibiotics) were collected, centrifuged, and resuspended in saline buffer. Ten microliters of each sample were used for a PCR with the following primers: Forward: GGCGAATGGGTGAGTAAC; Reverse: CGGATAACGCTTGCGACCT. Only negative samples were used in the study.

### Immunofluorescence staining

Cortical organoids were fixed with 4% paraformaldehyde overnight at 4°C and then transferred to 30% sucrose. After the 3D structures sink, they were embedded in O.C.T. (Sakura, Tokyo, Japan) and sliced in a cryostat (20 μm slices). Following air dry, the slides containing the sliced samples were permeabilized/blocked with 0.1% triton X-100 and 3% FBS in PBS for 2 hours at room temperature, and incubated with primary antibodies overnight at 4°C. Primary antibodies used in this study were: mouse anti-Nestin, Abcam (Cambridge, UK) ab22035, 1:250; rat anti-CTIP2, Abcam ab18465, 1:500; rabbit anti-SATB2, Abcam ab34735, 1:200; chicken anti-MAP2, Abcam ab5392, 1:2000; rabbit anti-Synapsin1, EMD-Millipore AB1543P, 1:500; mouse anti-NeuN, EMD-Millipore MAB377, 1:500; rabbit anti-Ki67, Abcam ab15580, 1:1000; rabbit anti-SOX2, Cell Signaling Technology 2748, 1:500; rabbit anti-GFAP, DAKO Z033429, 1:1000; rabbit anti-TBR1, Abcam ab31940, 1:500; rabbit anti-TBR2, Abcam ab23345, 1:500; rabbit anti-beta-catenin, Abcam E247, 1:200; mouse anti-GABA, Abcam ab86186, 1:200; rabbit anti-PROX1, Abcam ab101651, 1:250. Next, the slices were washed with PBS and incubated with secondary antibodies (Alexa Fluor 488-, 555- and 647-conjugated antibodies, Life Technologies, 1:1000) for 2 hours at room temperature. The nuclei were stained using DAPI solution (1 μg/mL). The slides were mounted using ProLong Gold antifade reagent and analyzed under a fluorescence microscope (Axio Observer Apotome, Zeiss).

### Synaptic puncta quantification

Pre-synaptic Syn1+ puncta were quantified after 3D reconstruction of z-stacks of random images from randomly selected regions of all lines and from two independent experiments. Only puncta overlapping MAP2-positive processes were scored.

### Electron microscopy (EM)

EM was performed at the CMM Electron Microscopy Facility at University of California San Diego. Four-month-old organoids were immersed in modified Karnovsky’s fixative (2.5% glutaraldehyde and 2% paraformaldehyde in 0.15 M sodium cacodylate buffer, pH 7.4) for at least 4 hours, post fixed in 1% osmium tetroxide in 0.15 M cacodylate buffer for 1 hour and stained in 2% uranyl acetate for 1 hour. Samples were dehydrated in ethanol, embedded in Durcupan epoxy resin (Sigma-Aldrich), sectioned at 50 to 60 nm on a Leica Ultracut UCT (Leica, Bannockburn, IL), and transfer onto Formvar and carbon-coated copper grids. Sections were stained with 2% uranyl acetate for 5 minutes and Sato’s lead stain for 1 minute. Grids were analyzed using a JEOL 1200EX II (JEOL, Peabody, MA) transmission electron microscope equipped with a Gatan digital camera (Gatan, Pleasanton, CA).

### Targeted single-cell qRT-PCR and analysis

Specific target amplification was performed in individual dissociated cortical organoids using C1 Single-Cell and BioMark HD Systems (Fluidigm, San Francisco, CA, USA), according to the manufacturer’s protocol and as previously described (Thanathom et al., 2016). Briefly, cortical organoids were mechanically dissociated after 30 minutes of incubation in Accumax (Innovative Cell Technologies, San Diego, CA, USA) at 37 °C under rotation. After passing through 100-μm and 40-μm strainers, cells were centrifuged and resuspended in Media2 (see *Generation of cortical organoids*). Single cortical cells were captured on a C1 medium chip and cell viability was assessed using a LIVE/DEAD Cell Viability/Cytotoxicity kit (Life Technologies). The targeted single-cell qPCR was performed using DELTAgene primer pairs in the 96.96 Dynamic Array IFC chip. The results were analyzed using Fluidigm Real-time PCR Analysis Software and Singular Analysis Toolset 3.0 (Fluidigm).

### 10X genomics single-cell and analysis

After organoid dissociation, single cells were processed through the Chromium Single Cell Gene Expression Solution using the Chromium Single Cell 3’ Gel Bead, Chip and Library Kits v2 (10X Genomics, Pleasanton) as per the manufacturer’s protocol. In brief, single cells were resuspended in 0.1% BSA in PBS. Five thousand cells were added to each channel with an average recovery rate of 1,746 cells. The cells were then partitioned into Gel Beads in Emulsion in the Chromium instrument, where cell lysis and barcoded reverse transcription of RNA occurred, followed by amplification, shearing and 5’ adaptor and sample index attachment. Libraries were sequenced on an Illumina HiSeq 2500. De-multiplexing, alignment to the hg19 transcriptome and unique molecular identifier (UMI)-collapsing were performed using the Cellranger toolkit (version 2.0.1) provided by 10X Genomics. A total of 3,491 cells with approximately 53,000 reads per cell were processed. Analysis of output digital gene expression matrices was performed using the Seurat R package. Matrices for replicates were merged with the MergeSeurat function and all genes that were not detected in at least 5% of all single cells were discarded, leaving 10,594 genes for further analyses. Cells with fewer than 600 or more than 8,000 expressed genes as well as cells with more than 50,000 UMIs or 0.1% mitochondrial expressed genes were removed from the analysis. Data were log normalized and scaled to 10,000 transcripts per cell. Variable genes were identified with the FindVariableGenes function. Principal components were evaluated for statistically significant gene expression signals using the JackStraw function. PCA was carried out, and the top 36 principal components were retained. With these principal components, t-SNE was applied with the RunTSNE function to visualize the cells in two dimensions and identified distinct cell clusters with the FindClusters function with resolution = 0.30. Differential expression to identify cluster markers was performed using the FindAllMarkers function.

### Data availability

All data and/or analyses generated during the current study are available from the corresponding author upon reasonable request. Single-cell RNA sequencing data that support the findings of this study have been deposited at NCBI GEO: GSE113089.

### Multi-electrode array (MEA) recording

Six-week-old cortical organoids were plated per well in 12-well MEA plates (Axion Biosystems, Atlanta, GA, USA). Each well contains 64 platinum microelectrodes with 30 μm of diameter spaced by 200 μm, yielding a total of 512 channels. The plate was previously coated with 100 μg/mL poly-L-ornithine and 10 μg/ml laminin, and we performed four independent experiments in duplicates. Cells were fed twice a week and measurements were collected 24 hours after the medium was changed, once a week, starting at two weeks after plating (8 weeks of organoid differentiation). Recordings were performed using a Maestro MEA system and AxIS Software Spontaneous Neural Configuration (Axion Biosystems) with a customized script for band-pass filter (0.1-Hz and 5-kHz cutoff frequencies). Spikes were detected with AxIS software using an adaptive threshold crossing set to 5.5 times the standard deviation of the estimated noise for each electrode (channel). The plate was first allowed to rest for three minutes in the Maestro device, and then four minutes of data were recorded. For the MEA analysis, the electrodes that detected at least 5 spikes/min were classified as active electrodes using Axion Biosystems’ Neural Metrics Tool. Bursts were identified in the data recorded from each individual electrode using an inter-spike interval (ISI) threshold requiring a minimum number of 5 spikes with a maximum ISI of 100 ms. A minimum of 10 spikes under the same ISI with a minimum of 25% active electrodes were required for network bursts in the well. The synchrony index was calculated using a cross-correlogram synchrony window of 20 ms. Bright-field images were captured to assess for cell density and electrode coverage.

### Custom MEA analysis

Custom MEA analysis and neonatal EEG/organoid LFP regression model can be found in: https://github.com/voytekresearch/OscillatoryOrganoids. Raw MEA recordings were converted to .*mat* files using Axion-provided functions and analyzed offline using custom MATLAB functions and scripts. Local field potential signals (LFP) from each of the 64 electrodes were generated by low-pass filtering (FIR filter) and downsampling raw signals from 12,500 Hz to 1,000 Hz (*resample.m*). Multi-unit spikes were detected as follows: each channel was first referenced to the well median for every time point, similar to a common average reference (64 channels). The median was used instead of the mean to avoid biasing the reference during high firing rate periods. Next, the re-referenced signal was bandpass filtered for 300-3,000 Hz with a 3rd-order Butterworth filter (*butter.m*). The spike threshold was set to be 5.5 standard deviations, where the standard deviation was estimated as previously described (Quiroga et al., 2004) to avoid biasing the threshold for channels with high firing rates (thus an artificially high threshold). Spike timestamps were taken as the peak time after the absolute value of the signal crossed the threshold, but at least 1 ms from another spike (*findpeaks.m*). Spike timestamps were then converted into binary vectors (1 ms bin size), summed across 64 channels, and smoothed (*conv.m*) with a normalized 100-point Gaussian window (*gausswin.m*) to create a population spiking vector for each MEA well. Note that spikes from each channel do not represent putative single-unit action potentials, as the spatial resolution of MEA electrodes were too sparse. Multi-unit spiking was not sorted since total population spiking (of well) was submitted for further analysis, rather than individual spike trains.

### Network event analysis

A network event was detected when population spiking was i) greater than 80% of the maximum spiking value over the length of the recording; ii) at least 1 spike/s; and iii) 1 second away from any other network events. The first peak after all 3 criteria was satisfied was marked as t = 0, and the window of data from 0.5 s before to 2.5 s after the peak was collected as the network event, as events almost always subsided 2.5 seconds after onset by both algorithmic detection and visual inspection. Nearly all spiking channels experienced a significant firing rate increase during network events. LFP data from all 64 channels from the same timeframe were also collected for analysis. All events from different MEA wells obtained on the same recording day were aggregated for statistical analysis and plotting. Subpeaks within an event were identified using *findpeaks.m*, where a subpeak must satisfy the following: i) peak height of at least 25% of the first peak; ii) peak width of at least 50 ms; iii) at least 200 ms away from the previous peak; and iv) peak prominence of 1 over Peak 1 height. Subpeak time and the height relative to the initial peak were recorded. The inter-event interval coefficient of variation (IEI CV) was calculated as the standard deviation of the inter-event interval divided by its mean, where IEI is the time between consecutive network events within the same MEA well. Event temporal correlation was calculated as the mean Pearson correlation coefficient of population spiking vector during each network event with every other network event in the same MEA well across a single recording session. Event spatial correlation was calculated as the mean Pearson correlation coefficient between all pairs of 64 LFP channels during each 3-s network event.

### Oscillatory spectral power analysis

Power spectral density (PSD) estimates were computed using Welch’s method (*pwelch.m*), with a window length of 2 s and overlap of 1 s. Oscillatory power was defined as peaks in the PSD above the aperiodic *1/f* power law decay. Thus, for each channel, a straight line was fit over the PSD in double-log space between 0.5-20 Hz using robust fit (*robustfit.m*), and oscillatory power was computed as the difference between the mean log PSD power and the mean log fitted power (baseline), over 2.5-4.5 Hz. This method accounts for non-oscillatory changes, such as slow transients or the aperiodic 1/f background component, whereas standard wavelet filtering methods will confound the two (Haller et al. 2018).

### Regression models

For analysis in Figure 2F, G and S7C, F, G, we fit regression models (*LinearModel.fit,MATLAB*) using organoid age (in days) as input and electrophysiological features as output. Order-1 (linear) models were fit for Figure 2G and and S7C, G, and order-2 (quadratic) models were fit for Figure 2F, 3C and Figure S7F. Reported R^2^ and p values are model statistics over the entire dataset. All events from different MEA wells on the same recording day were aggregated as samples drawn from the same distribution. To estimate culture age, we used 3 electrophysiological features as input: event latency, event peak spiking, and oscillatory power; and their square roots to account for the nonlinear inverted-U features. These were used to build a regression model. Within-well models were fit over all data points of the same well, and goodness-of-fit was reported as the model R^2^ and the RMSE. Across-well models were trained and evaluated using leave-1-out cross-validation, and goodness-of-fit is reported as the R^2^ and the RMSE computed over the validation set, not the training set.

### Phase Amplitude Coupling (PAC)

LFP data from all 64 channels of each well was first lowpass/bandpass filtered (*eegfilt.m*, EEGLAB) for delta (0-4 Hz) and high-frequency, broadband (100-400 Hz) activity, sometimes referred to as high gamma. Delta phase was extracted by taking the phase angle of the bandpassed delta signal Hilbert transform (*hilbert.m, angle.m*), while gamma power was extracted by taking the squared magnitude of the filtered gamma. Gamma power was smoothed with the same delta-band filter for display purposes, but not for subsequent analysis. Note that analysis was performed for 100-200 Hz and 200-400 Hz separately, as LFP spectrum follows an inverse power law (1/f), and grouping a wide frequency band (100-400 Hz) together would bias power estimates towards lower frequency limits (∼100 Hz). To compute PAC, instantaneous delta phase was binned into 20 equidistant bins between -π and π, and gamma power was sorted based on the corresponding delta phase at the same sample time and averaged across the same phase bin. This procedure was performed separately for event and non-event indices, where event indices are the same 3-second windows as described above in *Network Event Analysis,* while all other times are considered as non-event time points. Modulation Index was computed as the Kullback-Leibler divergence between the sum-normalized distribution of gamma power across phase bins and a uniform distribution (Tort et al., 2010). Figure 3C presents well-averaged MI across all 64 channels. For visualization in Figure 3B, the binned gamma vector for each channel was circularly shifted such that the phase of maximum gamma power was -π.

### Pharmacology

The pharmacological manipulation was performed using the following drugs: 10 μM bicuculline, 50 μM muscimol, 20 μM CNQX, 20 μM AP5, 25 μM baclofen and 1 μM TTX. In this assessment, baseline recordings were obtained immediately before and 15 minutes after the addition of the compound. Three washes with PBS for total removal of the drug were performed in washout experiments; fresh media was added and another recording was conducted after 2 hours.

### Preterm neonatal EEG

A preterm neonatal EEG dataset was obtained elsewhere (Stevenson et al., 2017). Raw recordings were not available due to patient confidentiality concerns. The dataset includes 567 recordings from 39 preterm neonates (24-38 weeks old conception age), consisting of 23 EEG features computed from the entirety of each recording and the post-conception age in weeks (Table S2).

### Neonate-organoid age correlation model

To compare the developmental trajectory of cortical organoids and the preterm human brain, we trained an Elastic Net (L1- and L2-regularized) regression model on only the preterm neonatal EEG features and used that model (with all parameters held the same) to generate an equivalent organoid “brain-age” for each recording time point over 40 weeks in culture. Specifically, the training dataset consisted of a subset of the preterm EEG data; we discarded all “low-activity-period” features (Lisman, 1997) since there was no equivalent period for organoid recordings, as well as features for which we could not sensibly compute from organoid LFPs, such as interhemispheric synchrony. This selection was done *a priori*, and 13 features remained, including 4 features for relative spectral power in distinct frequency bands, which were further discarded due to frequency-dependent filtering properties of the skull and difference in spatial integration of currents in macroscopic EEG electrodes compared to microscopic planar MEA electrodes. The remaining 9 features correspond to aspects of spontaneous activity transient (SAT) timing, such as SATs per hour and SAT duration, which were similarly computed on organoid LFPs after network event detection described earlier (see Table S2 for a full list of included and rejected features). This latter organoid LFP test dataset was never seen by the regression model until prediction time. Training was performed using scikit-learn linear model module [(ElasticNetCV (Pedregosa et al., 2011)], with K-Group shuffle split cross-validation on regularization hyperparameters, where K = 25% of groups, N = 200 shuffles. In other words, we found the best regularized linear model possible for predicting the conception age of preterm neonates using those 9 precomputed EEG features. This model was directly applied on organoid LFP features to determine the corresponding “brain age” of the organoids during 40 weeks in culture. 1-sample *t*-tests were performed from every time point to test whether the mean estimated “brain age” was significantly different from the organoid culture age.

### Resampled feature correlation

We computed Pearson’s correlation coefficient between neonate age and each of the 9 EEG features, after a leave-K-groups-out resampling procedure N times, where K is the number of neonates from whom all recordings were left out in computing the correlation (50% of all neonates, resampling N = 100). An identical procedure was performed to compute the correlation between organoid culture age and LFP features (K = 4 out of 8, 50%, N = 100). Mean and standard deviation were then computed over all resampled draws in order to compare between organoid LFP and neonatal EEG.

### Statistical analysis

Data are presented as mean ± s.e.m., unless otherwise indicated, and it was obtained from different samples. No statistical method was used to predetermine the sample size, and no adjustments were made for multiple comparisons. The statistical analyses were performed using Prism software (GraphPad, San Diego, CA, USA). Student’s *t*-test, Mann–Whitney-test, or ANOVA with post hoc tests were used as indicated. Significance was defined as P < 0.05(*), P < 0.01(**), or P < 0.001(***). Blinding was used for comparing affected and control samples.

## SUPPLEMENTAL FIGURE LEGENDS

**Figure S1. Cellular and molecular characterization of human cortical organoids.** (A) Schematic of the protocol used to generate cortical organoids. Scale bar, 200 μm. (B) Reproducibility of organoid size at 2 months of maturation (*n = 20* independent experiment, 7 different cell lines). (C) Organoids are composed of a proliferative region surrounded by intermediate progenitor cells, cortical and GABA+ neurons. Scale bar, 50 μm. (D) Principal component analysis (PCA) of cells projected onto the first two components. Overlaid populations of 2- and 10-month-old cortical organoids are compared to 2-month-old 2D monolayer neurons. All timelines for this and the subsequent experiments consider the iPSC stage as day 0 (*n = 2* independent cell lines for each cortical culture; *n = 3* for 2D monolayer neurons). (E-F) Violin plots illustrate the differences in single-cell expression of target genes in cortical organoids and 2D neurons. (G-H) Unsupervised hierarchical clustering single-cell analysis. Genes were clustered using the Pearson correlation method and cells were clustered using the Euclidean method.

**Figure S2. Reproducibility and cell diversity in cortical organoids.** (A) Schematic showing the single-cell approach performed to access reproducibility of organoid generation using two control iPSC lines. (B) *t*SNE plot of single-cell mRNA sequencing data from 6-month-old organoids color-coded by replicate. (C) Split Dot Plot depicting the correlation between expression patterns of representative markers and cell populations identified within the dataset. The size of the dots represents the percentage of cells expressing a given gene, while the intensity of the color denotes the average expression level (grey, low expression; red/blue, high expression). (D) Population ratio of each cluster by replicate. (E) Violin and *t*SNE plots of selected genes depicting the proportion of cells contributing to each cluster. For the violin pots, the dot denotes a cell while colors correspond to their cluster identity. (F) The *t*SNE plots show the contribution of an individual cell-type marker within each cluster (red denotes higher expression).

**Figure S3. Long-term MEA network activity.** (A) Representative bright-field image of cortical organoids on the MEA plate. (B) Schematic representation of the electrical activity features analyzed from the MEA recordings. Each bar represents a spike; and a spike cluster (in blue) represents a burst. Bursts occurring at the same time in different channels characterize a network burst. The synchrony index is based on the cross-correlogram and represents a measure of similarity between two spike trains. (C) Temporal evolution of network activity characterized by different parameters. (D) Raster plots illustrating the development of network activity. (E) Consistent and reproducible development of electrical activity in cortical organoids over time. The data are shown as mean ± s.e.m (*n = 8*, independent experiments performed in duplicates using two clones of a control iPSC line).

**Figure S4. Extended characterization of network electrophysiology.** (A) Spikes detected on 9 channels. Black traces represent single spikes, blue and red traces represent the average of positive and negative spikes, respectively. Spike trains are not sorted for their polarity in the subsequent analyses, as total population spiking is the main feature of interest. (B) Representative oscillatory network events. Each overlapping trace represents a single occurrence of an event recorded on the same channel. LFP polarity of events differs between channels due to the spatial configuration of cells around the electrode. (C) Event onset peak (Peak 1) increases in amplitude until 30 weeks, while (D) subpeak amplitude continues to increase (for the 2nd-4th peak) throughout development. (E) Subsequent peaks occur with a consistent latency of ∼400 ms after the previous peak, particularly for Peak 3 and 4. (F) Temporal similarity of network events during the 3-s window is high at early time points, but decreases with development, acquiring more variable dynamics within an event. (G) Temporal similarity of network events during the 3-s window is high at early time points, but decreases with development, acquiring more variable dynamics within an event. The data showed in C, F and G are presented as mean ± s.e.m., linear (C, G) or quadratic (F) model regression. (H) Comparison of the protocol for neurosphere and cortical organoid generation. (I) Network-wide giant depolarizing potentials occur at a similar rate to those found in organoids recordings, and visible perturbations are observed in the LFP trace. However, the network recruitment in neurospheres is lower than in organoids (less than 8 spikes/s), and events have significantly shorter duration. No coherent low-frequency depolarizations are observed in filtered LFP events (J).

**Figure S5. Network activity in cortical organoids resembles oscillatory features in the developing human brain.** (A) Spectral representation of time series data from a 6-month-old cortical organoid, demonstrating oscillatory phenomenon. Spectrogram (left) of organoid LFP shows bursts of activity localized at low frequencies, while power spectral density (PSD, right) displays canonical *1/f* power law decay and a narrow oscillatory peak at 3 Hz. (B) Comparison of 9 preterm neonate EEG and cortical organoid features over time. For included EEG features, see Table S2. (C) Distributions of resampled Pearson correlation coefficients between feature and age for preterm neonate and organoid.

## SUPPLEMENTAL TABLES

**Table S1.**
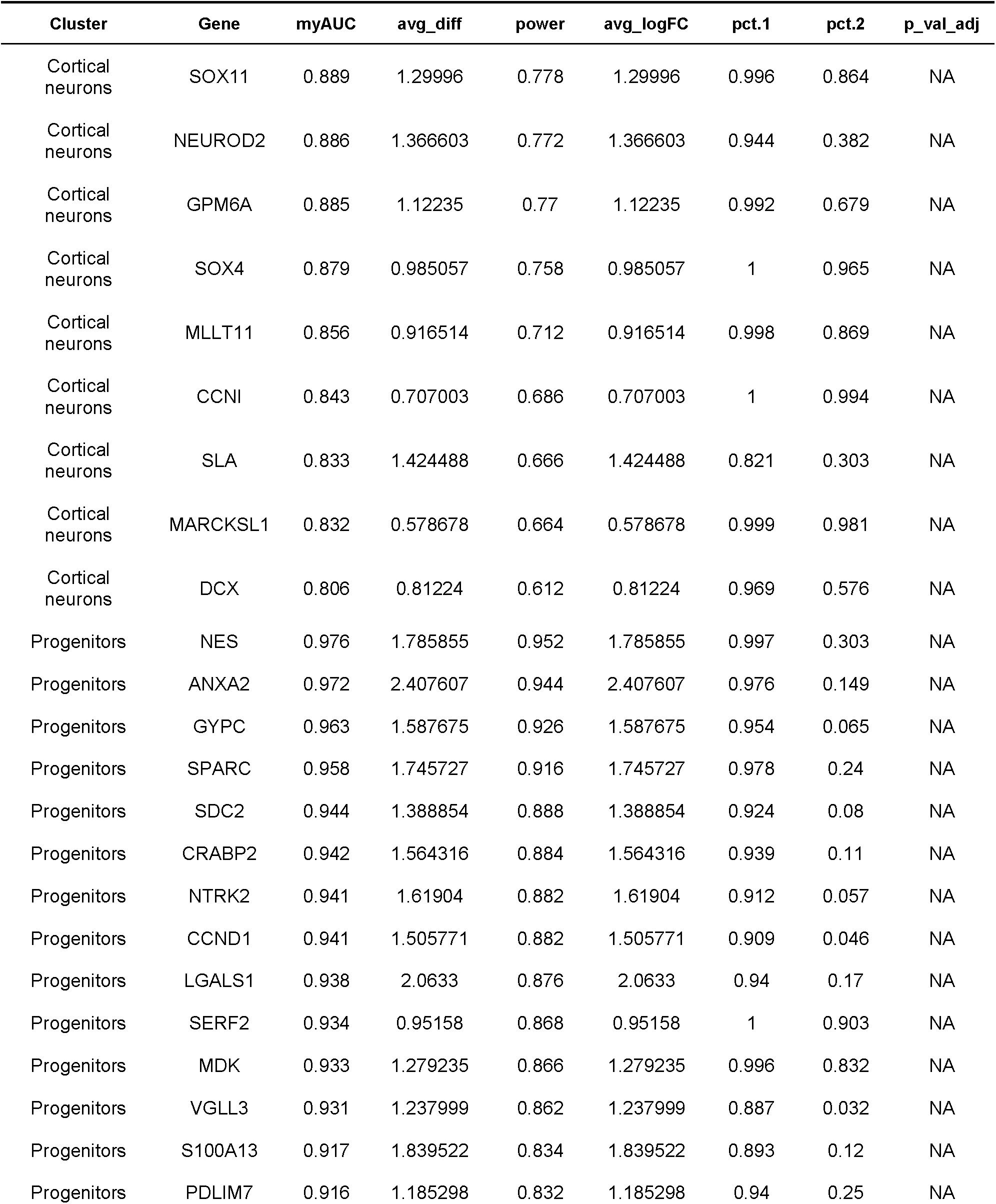
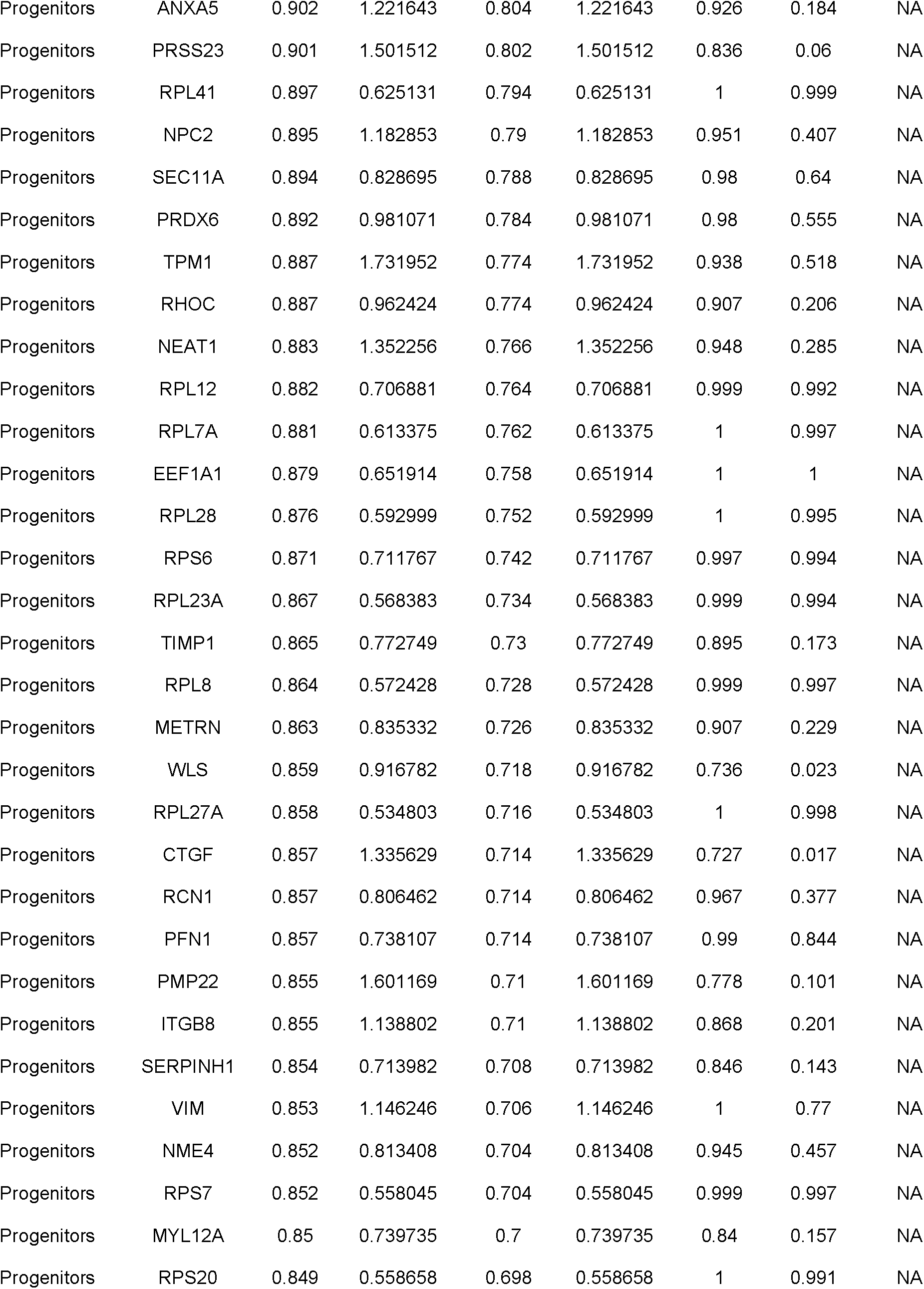
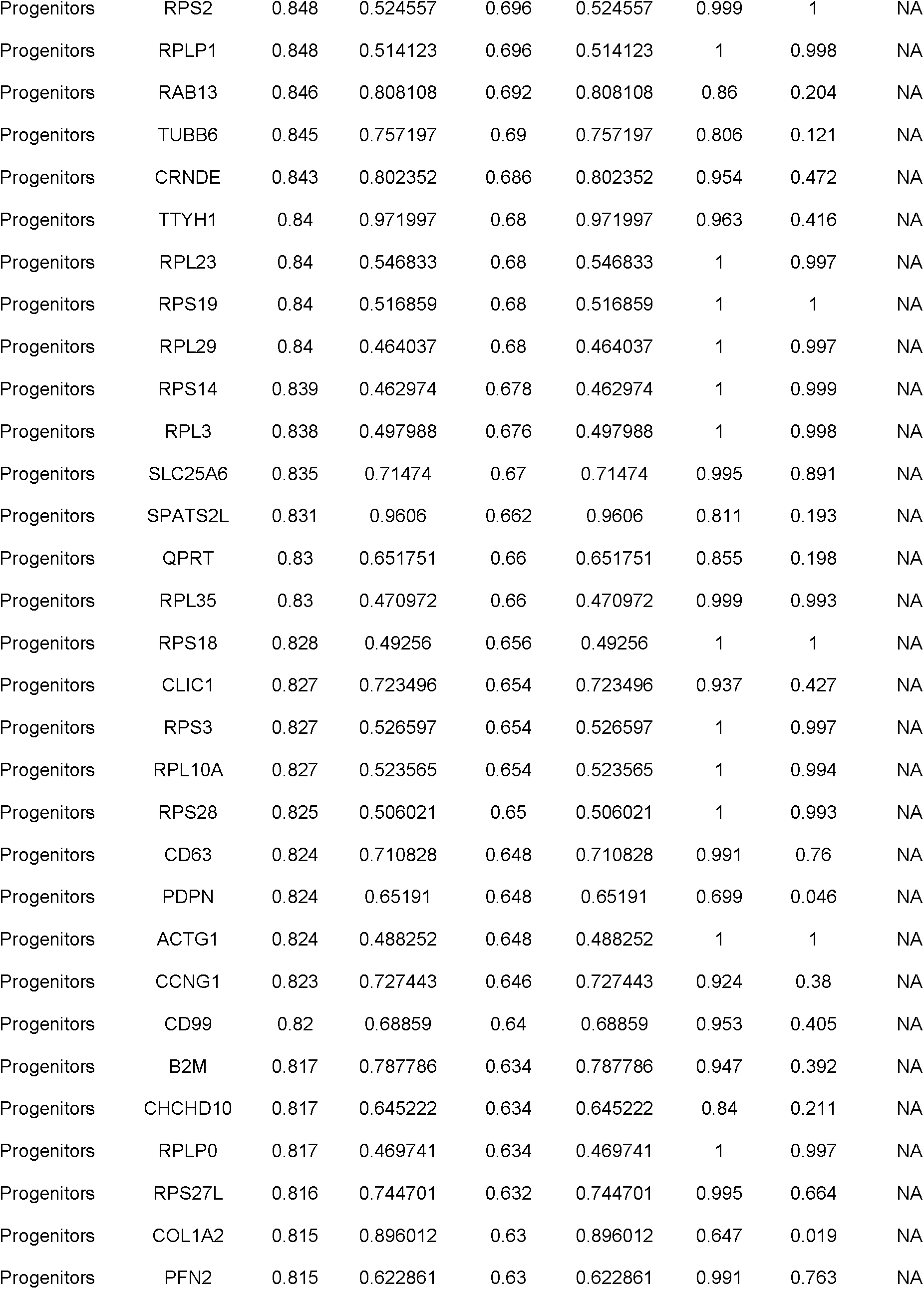
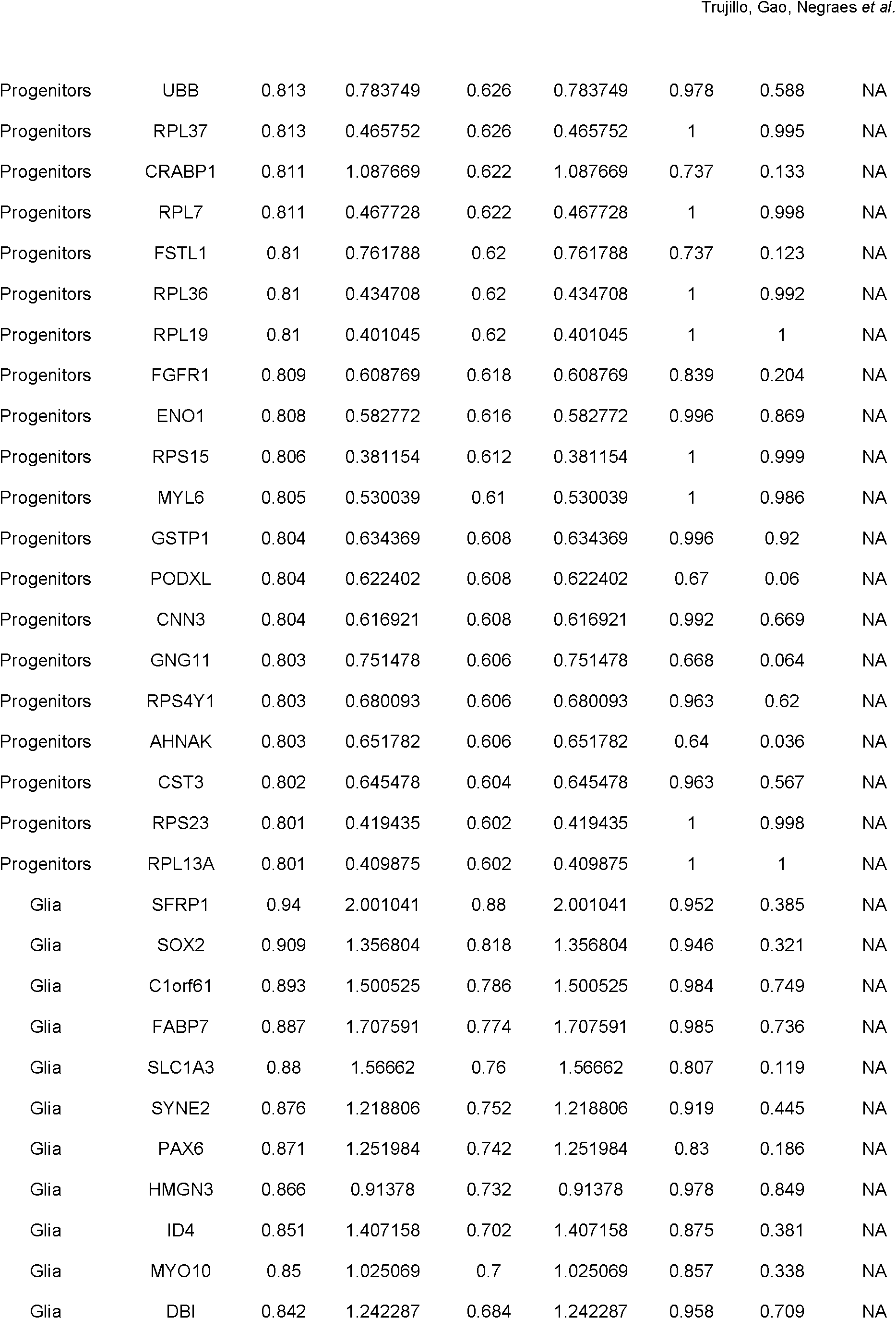
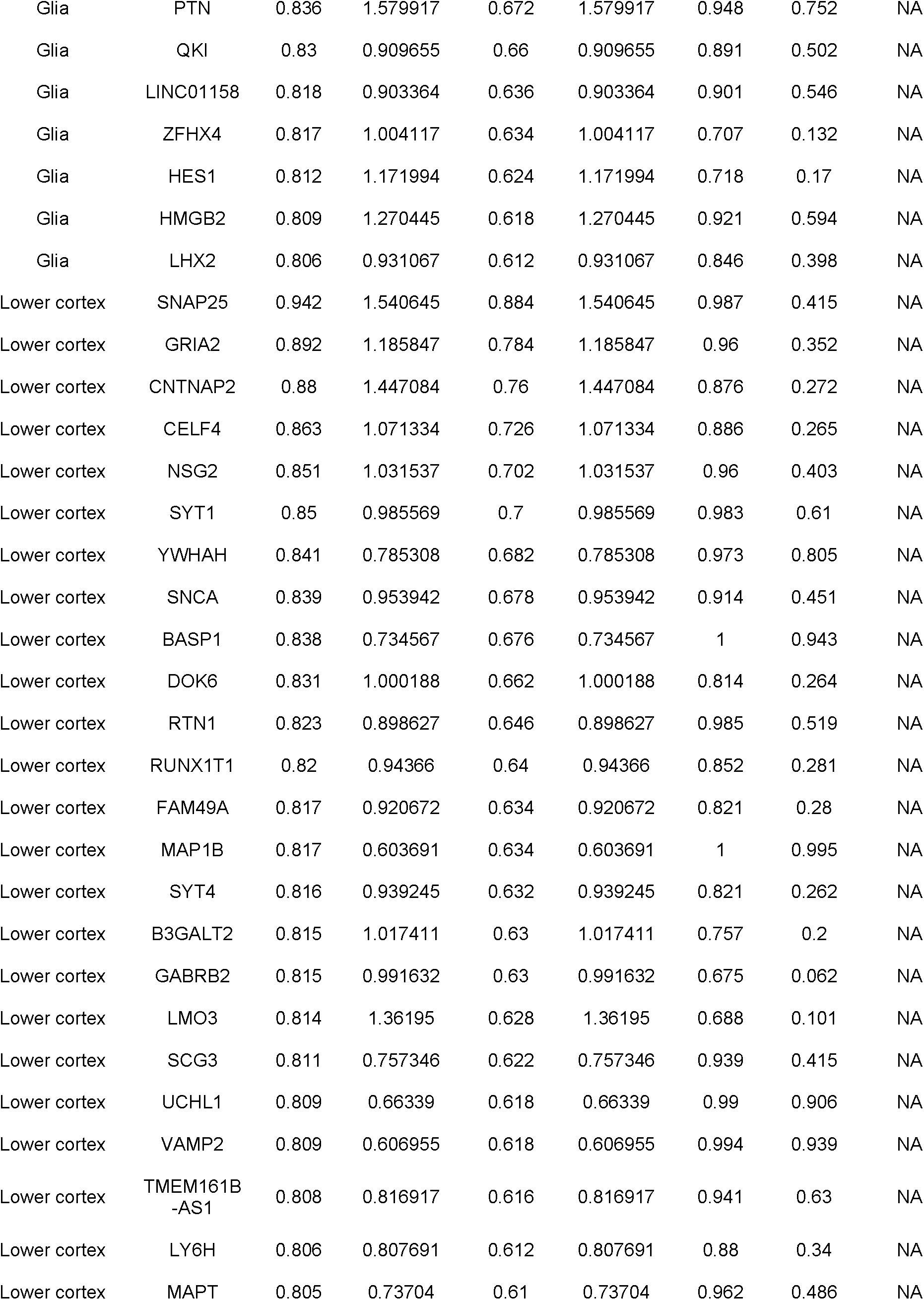
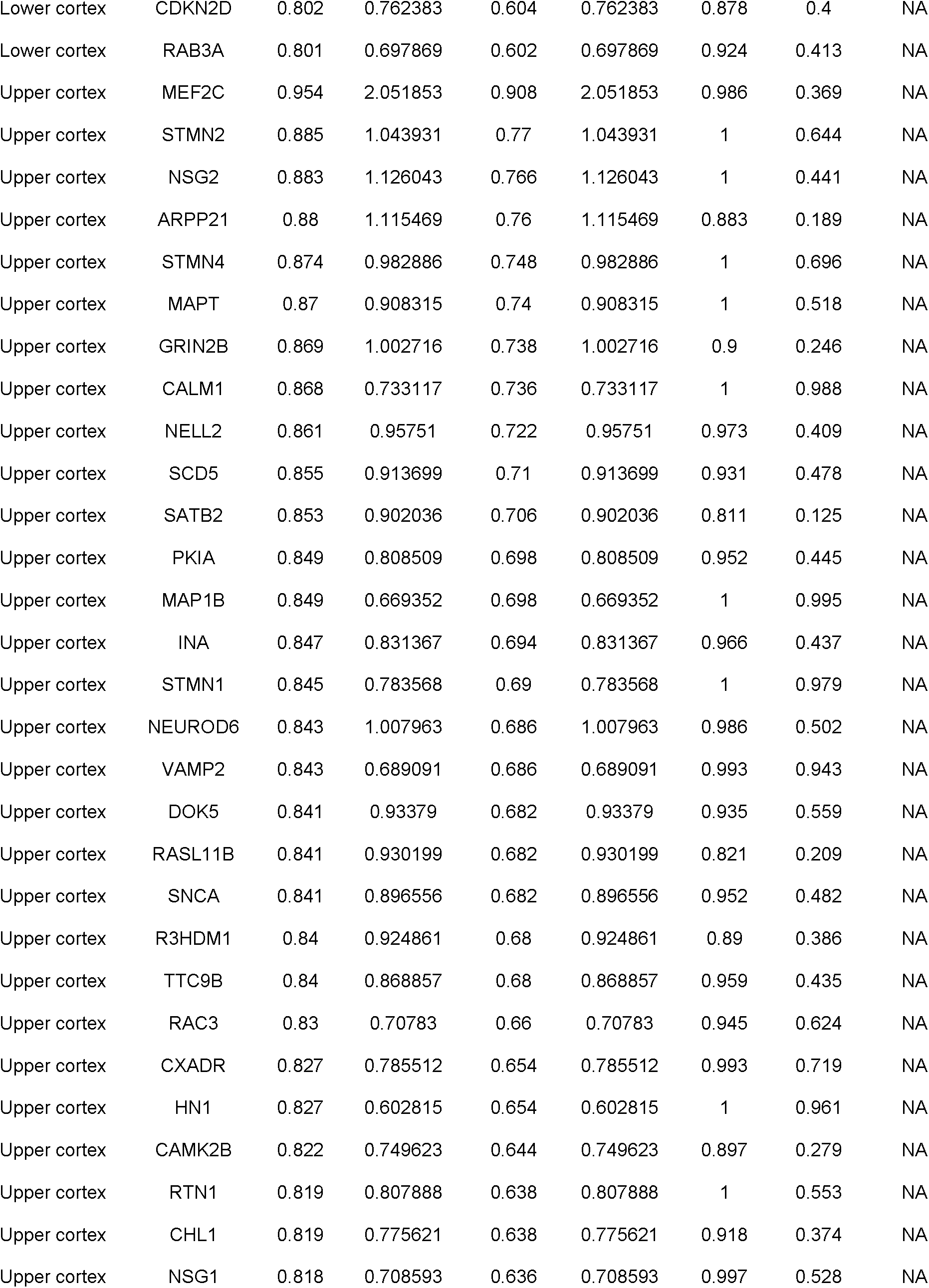
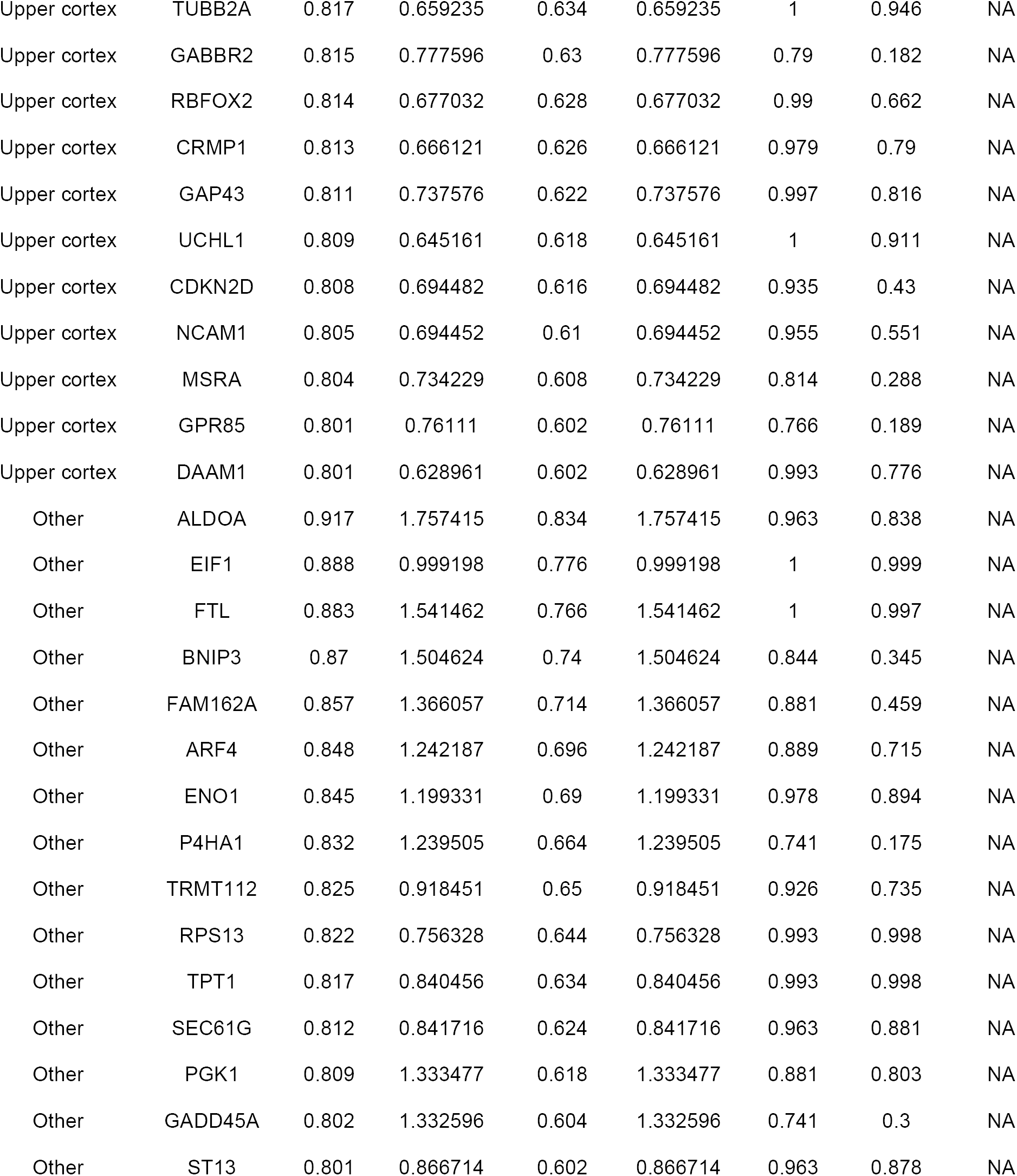
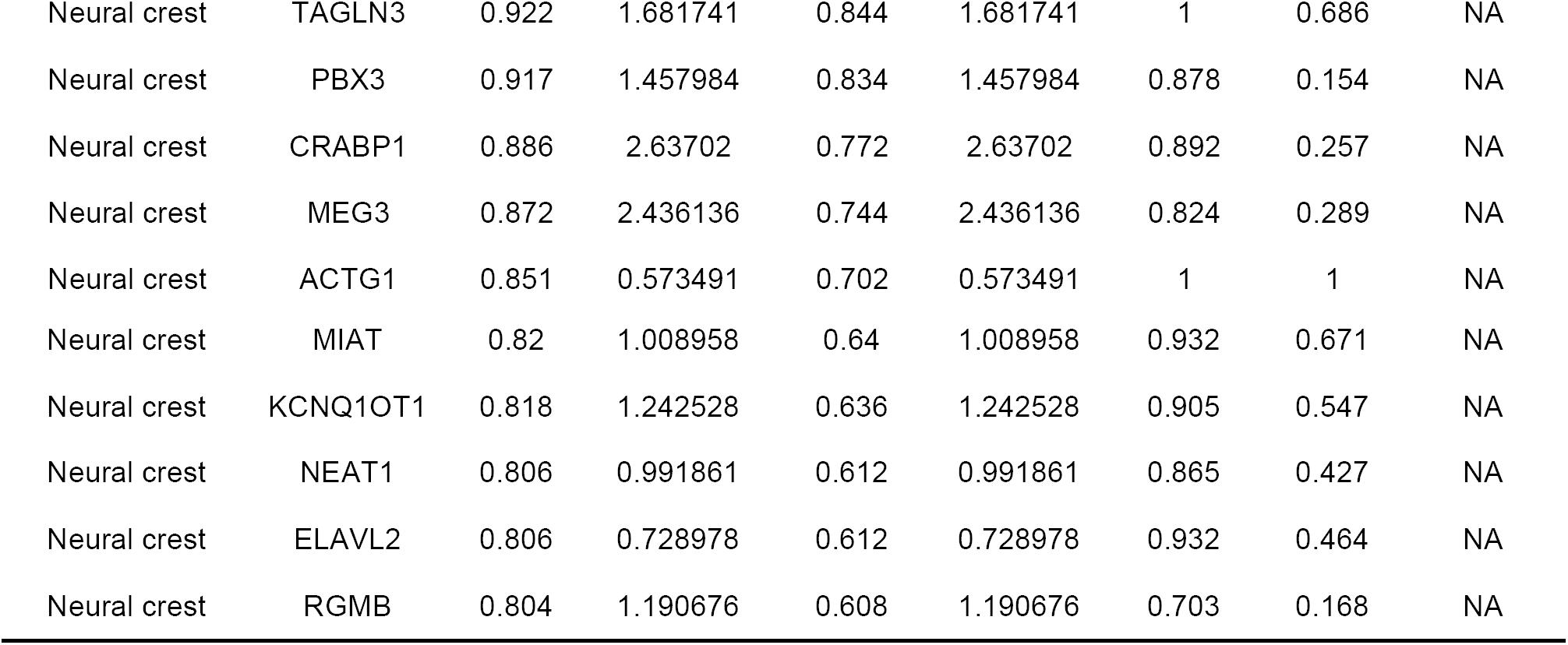
Top expressed genes of each cell cluster.

**Table S2.**
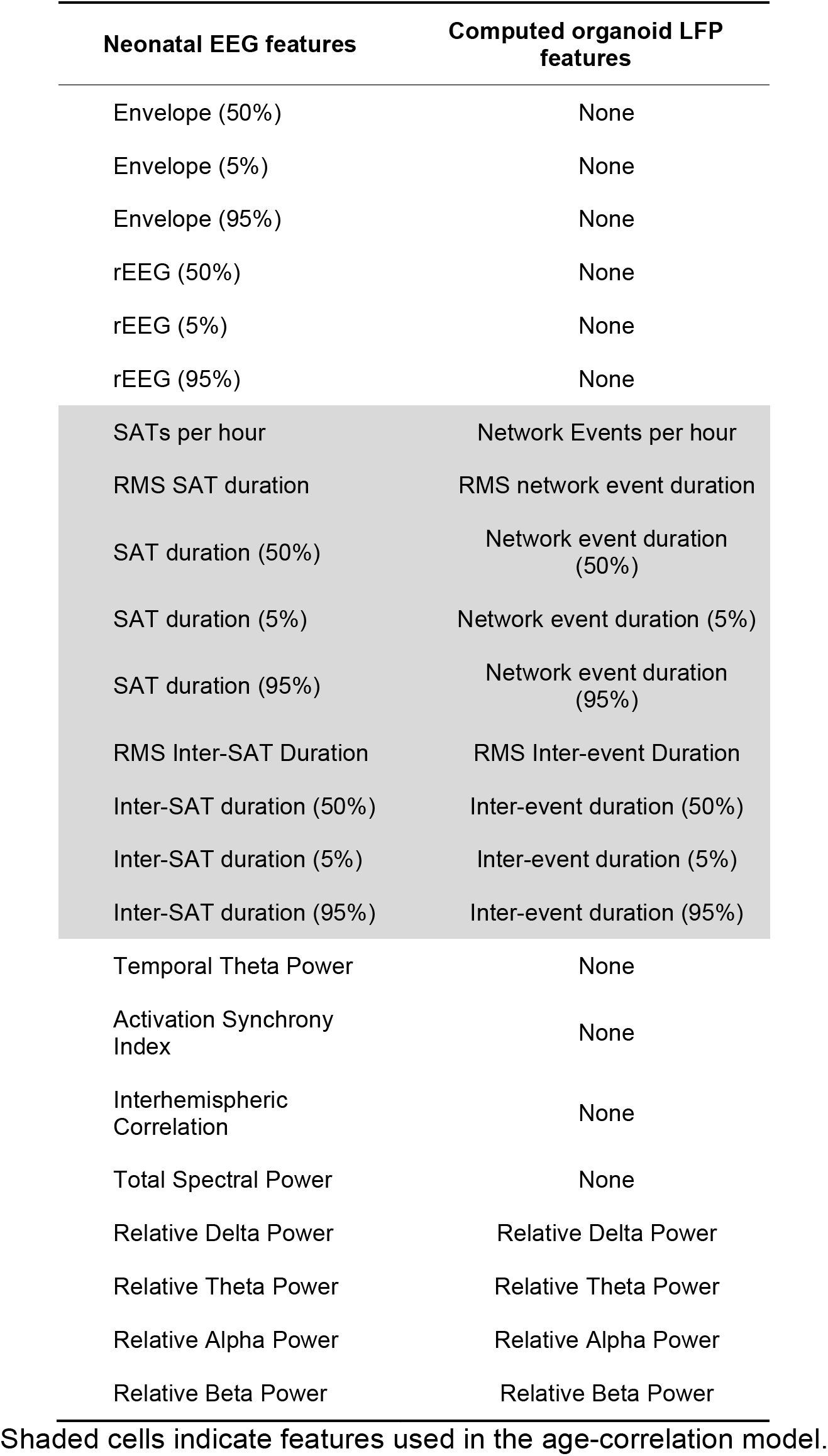
Electrophysiological features in preterm neonatal EEG dataset and analogous features computed in organoid LFP.

| Neonatal EEG features | Computed organoid LFP features |
| --- | --- |
| Envelope (50%) | None |
| Envelope (5%) | None |
| Envelope (95%) | None |
| rEEG (50%) | None |
| rEEG (5%) | None |
| rEEG (95%) | None |
| SATs per hour | Network Events per hour |
| RMS SAT duration | RMS network event duration |
| SAT duration (50%) | Network event duration (50%) |
| SAT duration (5%) | Network event duration (5%) |
| SAT duration (95%) | Network event duration (95%) |
| RMS Inter-SAT Duration | RMS Inter-event Duration |
| Inter-SAT duration (50%) | Inter-event duration (50%) |
| Inter-SAT duration (5%) | Inter-event duration (5%) |
| Inter-SAT duration (95%) | Inter-event duration (95%) |
| Temporal Theta Power | None |
| Activation Synchrony Index | None |
| Interhemispheric Correlation | None |
| Total Spectral Power | None |
| Relative Delta Power | Relative Delta Power |
| Relative Theta Power | Relative Theta Power |
| Relative Alpha Power | Relative Alpha Power |
| Relative Beta Power | Relative Beta Power |

Shaded cells indicate features used in the age-correlation model.

